# Anti-phasic oscillatory development for speech and noise processing in cochlear implanted toddlers

**DOI:** 10.1101/2022.03.07.483211

**Authors:** Meiyun Wu, Yuyang Wang, Xue Zhao, Tianyu Xin, Kun Wu, Haotian Liu, Shinan Wu, Min Liu, Xiaoke Chai, Jinhong Li, Chaogang Wei, Chaozhe Zhu, Yuhe Liu, Yu-Xuan Zhang

**Affiliations:** State Key Laboratory of Cognitive Neuroscience and Learning, Beijing Normal University, Beijing 100875, China; Department of Otolaryngology Head and Neck Surgery, Beijing Friendship Hospital, Capital Medical University, Beijing 100050, China; Department of Otolaryngology Head and Neck Surgery, Hunan Provincial People’s Hospital (First Affiliated Hospital of Hunan Normal University), Changsha 410005, China; Department of Otolaryngology Head and Neck Surgery, West China Hospital of Sichuan University, Chengdu 610044, China; Department of Otolaryngology Head and Neck Surgery, Peking University First Hospital, Beijing 100034, China

**Author notes:** These authors contributed equally (Meiyun Wu, Yuyang Wang). **Correspondence:** (Y.Z.) and (Y.L.).

**Keywords:** cochlear implant, brain development, fNIRS, brain imaging, developmental trajectory, sound recognition, speech perception

## Abstract

Human brain demonstrates amazing readiness for speech and language learning at birth, but the auditory development preceding such readiness remains unknown. Cochlear implanted (CI) children with prelingual deafness provide a unique opportunity to study this developmental stage. Using functional near-infrared spectroscopy, we revealed that the brain of CI children was nearly irresponsive to sounds at CI hearing onset. With increasing CI experiences up to 32 months, the brain demonstrated function, region and hemisphere specific development. Most strikingly, the left anterior temporal lobe showed an oscillatory trajectory, changing in opposite phases for speech and noise. In addition, speech responses increased linearly in left sylvian parieto-temporal area and right inferior frontal gyrus, and noise responses changed in U shape in right supramarginal gyrus. Such cortical development predicted behavioral improvement. The study provides the first longitudinal brain imaging evidence for early auditory development preceding speech acquisition in the human brain.

**Graphic abstract:** 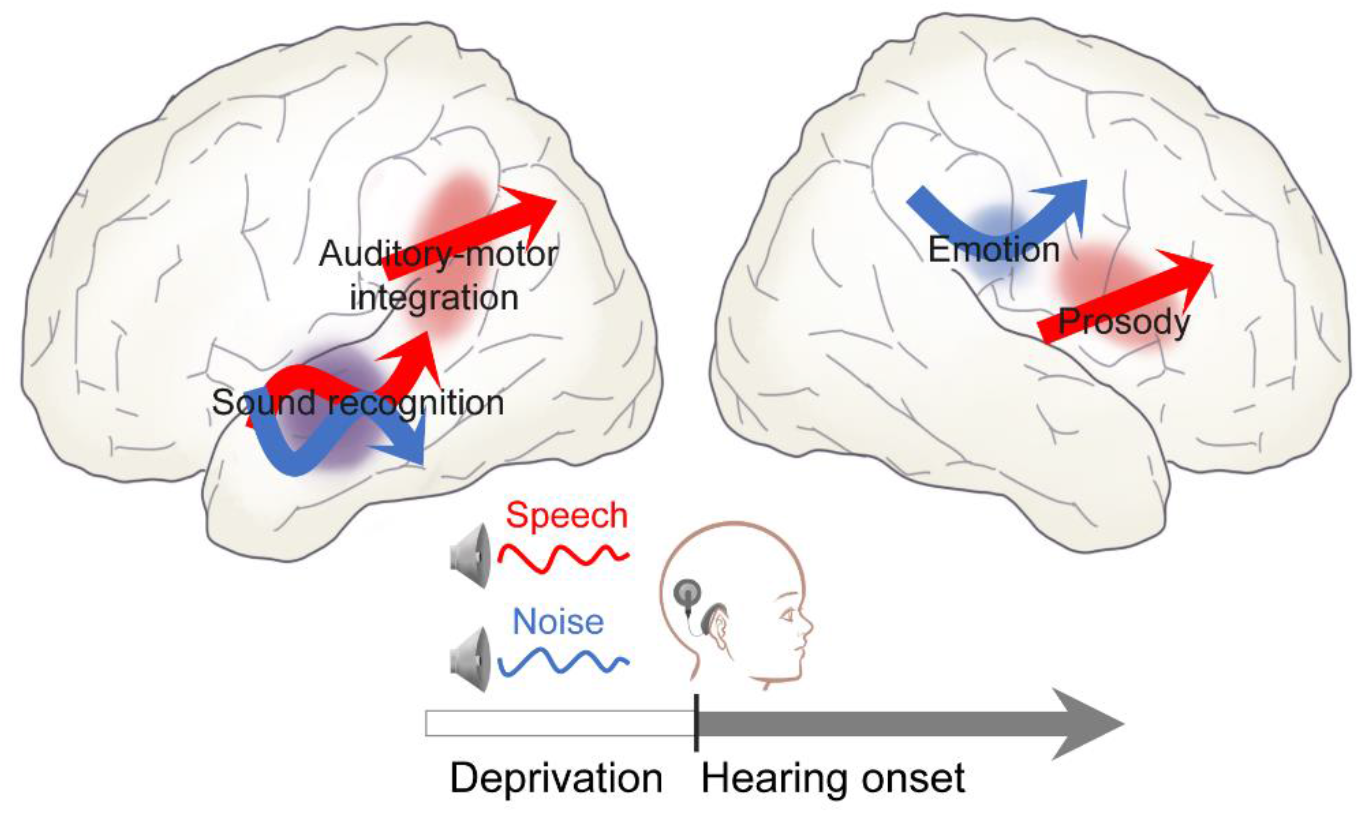

**Highlights:**

- The brain of cochlear implanted toddlers is functionally inferior to neonates
- Hearing experiences drive functional development in auditory related cortical areas
- Speech and noise processing develop in anti-phasic oscillatory trajectories
- Auditory cortical development predicts post-implantation behavioral improvement

## INTRODUCTION

Studies with infants revealed amazing readiness of the brain to undertake cognitive tasks such as speech and language learning at birth. For instance, resting-state functional magnetic resonance imaging revealed that preterm infants have shown similar functional networks to the adult brain (Doria, Beckmann et al. 2010, Turk, Heuvel et al. 2019). Longitudinal brain imaging showed that the auditory and sensorimotor networks developed the earliest, changing little during the first years of life (Gao, Wei et al. 2015). Indeed, consistent with the early onset of auditory function at approximately 24-26 gestational weeks (Birnholz and Benacerraf 1983, Eggermont and Moore 2012), the brain of fetuses and early preterm infants can respond to sounds (Schnupp, Nelken et al. 2011, Ghio, Cara et al. 2021) and discriminate phonetic contrasts in the bilateral superior temporal gyrus and inferior frontal gyrus (Mahmoudzadeh, Dehaene-Lambertz et al. 2013). However, due to the difficulty of measuring brain functions before birth, the process leading to such functional readiness of the auditory brain is unknown.

Auditory deprivation early in life delays cortical synaptogenesis, alters synaptic growth and pruning, myelination, and prevents maturation of auditory functions (Kral, Hartmann et al. 2001, Kral, Hartmann et al. 2002, Kral, Hartmann et al. 2002, Kral and A. 2013, Kral, Dorman et al. 2019). Hearing restoration via cochlear implant (CI) causes rapid synaptic growth to a higher level of synaptic connections within a short period and exaggerated pruning with auditory experience (Kral, Tillein et al. 2006, Kral and A. 2013, Kral, Dorman et al. 2019). The same developmental catch up is believed to occur in cochlear implanted children with prelingual severe sensorineural hearing loss, enabling them to acquire verbal language and communication (Kral, Dorman et al. 2019). Therefore, CI children provide a unique opportunity to study early auditory development of the human brain preceding speech acquisition.

Towards this end, we examined the functional development of auditory and speech related cortical networks in 67 young CI children (mean age 2.77 year ± 1.31 std) in a longitudinal brain imaging study using functional near-infrared spectroscopy (fNIRS). The fNIRS measures the hemodynamic changes in cortical surface via optical absorption properties of blood hemoglobin (Jöbsis 1977), with the advantage of presenting little noise, being compatible with the CI device, and friendly to young children, particularly for repeated measurements (Lu, Zhang et al. 2010, Sevy, Bortfeld et al. 2010, Cai, Dong et al. 2018). Post implantation performance on auditory perception and verbal communication were measured using clinical questionnaires with parents/guardians. Cortical imaging and performance measurements were conducted on each of the children’s hospital visits up to 32 months after CI activation. As a control for mature brain functions, 35 young healthy adults received fNIRS using the same protocol as the CI children.

## RESULTS

### Sound Evoked Cortical Responses at CI activation

To examine how the auditory and speech processing networks were affected by auditory deprivation early in life, we recorded sound evoked hemodynamic responses of implanted children using fNIRS on the day of their CI activation (N=30; see Figure S1A for testing schedule and table S3 for demographic detail), when the children were not yet exposed to electro-acoustic stimulation (Figure 1). As a control, 35 normal hearing young adults (NH adults) were also scanned using the same protocol. The oxy-hemoglobin (HbO) and deoxy-hemoglobin (Hb) changes showed the typical hemodynamic relationship for both CI children and NH adults during speech and noise stimulation (Figure 2A): HbO increased and Hb decreased shortly after auditory stimulation, with a time course similar to previous studies (for example, Mahmoudzadeh, Dehaene-Lambertz et al. 2013). Only averaged HbO amplitude during stimulation window was used for the following analyses, as it is known to be more strongly correlated with cognitive activities (Malonek, Dirnagl et al. 1997, Strangman, Culver et al. 2002, Fu, Mondloch et al. 2014) and more sensitive to the regional cerebral blood flow changes (Hoshi 2007). Linear mixed effect models (LME) allowing for separation of subject-level and group-level variations (Wierenga, Van et al. 2017, Bulgarelli, Klerk et al. 2020, Smith, Condy et al. 2020) were conducted.

**Figure 1.**
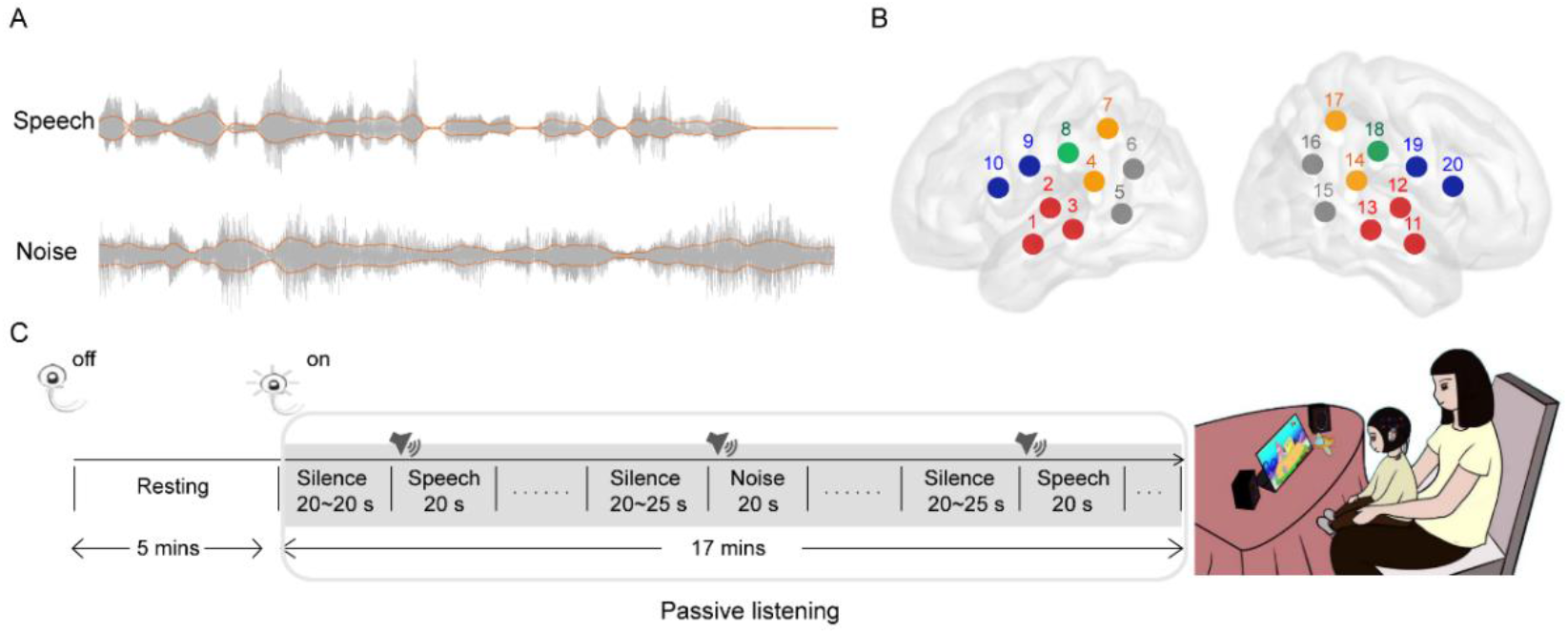
Experimental Design. (A) Sound waveforms of the speech and noise stimuli presented during functional near-infrared spectroscopy (fNIRS) recording, with amplitude envelopes, the primary acoustic signal carried by cochlear implants, highlighted in red lines. (B) Arrangement of the fNIRS channels, 10 on each hemisphere, localized using a transcranial brain atlas for children. The channels covered cortical areas related to auditory and language processing, including bilateral anterior temporal lobes (ATLs, in red), Sylvian parieto-temporal areas (Spts, in orange), supramarginal gyri (SMGs, in green), inferior frontal gyri (IFGs, in blue), and temporo-occipital junctions (in gray). (C) The fNIRS recording protocol. Cochlear implants and hearing aids if in wearing were turned off for the first 5 minutes of resting state, and then on for task-free sound presentation during the following 17 minutes. Sound conditions were presented from two speakers placed 45° left and right to the direction that the participant was facing in blocks of 20 seconds, separated by silence intervals of 20 to 25 seconds.

**Figure 2.**
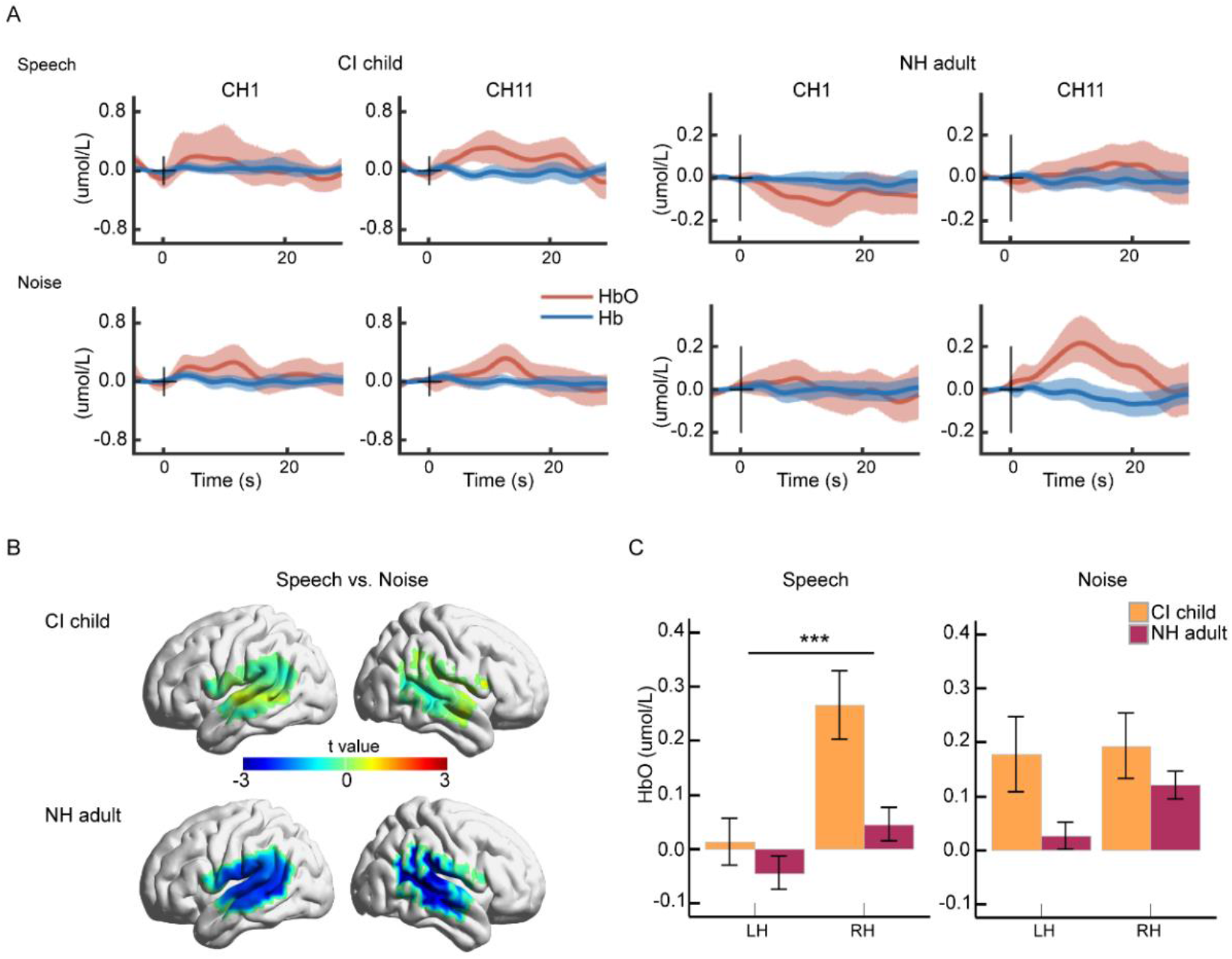
Cortical responses to sounds upon cochlear implant activation. (A) The oxy-hemoglobin (HbO, red) and deoxy-hemoglobin (Hb, blue) concentration changes for cochlear implanted children (CI child, N=30, left two columns) and normal hearing adults (NH adult, N=35, right two columns) in response to speech (top row) and noise (bottom row) in channel 1 and 11. The solid lines and the shaded areas represent mean and 95% confidence intervals calculated using bootstrapping (R=10,000). (B) T statistics map for speech and noise evoked HbO responses in CI children (top row) and NH adults (bottom row). (C) Comparison of CI child (orange) and NH adult (marron) HbO responses in the two channels showing sound evoked activity in CI children (channel 11 and 13 on the right anterior temporal lobe) and their counterparts on the left hemisphere (channel 1 and 3). Abbreviations: CI, cochlear implant; NH, normal hearing; LH, left hemisphere; RH, right hemisphere. Error bars indicate standard error of mean; ***p < 0.001.

Compared with silence, CI children showed no functional responses to sounds except for two channels in the right temporal lobe (p < 0.05, FDR corrected; channel 11 & 13, see Figure S1B), and little discrimination between speech and noise (Figure 2B, top row). In contrast, NH adults showed significant speech-noise discrimination not only in the temporal lobe but also in the frontal and parietal parts of the speech processing network (Figure 2B, bottom row). Furthermore, HbO responses in the two sound-activated channels for CI children were compared with NH adults for both sound conditions and hemispheres, with group, sound condition, and hemisphere as fixed factors using an LME model. Auditory responses were greater in magnitude in the right than in the left temporal cortex (Figure 2C; LME: main effect of hemisphere: *F*_(1,183.2)_ = 13.46, p < 0.001), in CI children than in NH adults (main effect of group: *F*_(1,64.3)_ = 7.18, p = 0.009), but showed only marginal overall difference between speech and noise (main effect of condition: *F*_(1,165.8)_ = 3.45, p = 0.065; all interactions: *F*s < 2.44, all p values > 0.120). Therefore, following auditory deprivation early in life, CI children showed higher cortical responsiveness to sounds than NH adults and lack of speech-noise discrimination, appropriate to an early stage of brain development.

### Longitudinal Development of Auditory Cortical Responses with CI Experiences

Longitudinal development of auditory cortical functions was examined in 57 CI children who completed at least 2 valid fNIRS assessments within an observation period up to 32 months (see table S3 for demographic detail). HbO responses often underwent non-monotonic changes with increasing electro-acoustic hearing experiences, particularly in the left temporal cortex (Figure 3A for a representative individual). Developmental trajectories were first evaluated in preliminary channel by channel analyses using LME models, with duration of CI experiences (CI_dur_) as main effect. Best models were selected based on the Akaike information criterion (AIC) and log-likelihood (logLik). For both speech and noise, the best LME models were cubic polynomial functions in the left temporal lobe (Figure 3B).

**Figure 3.**
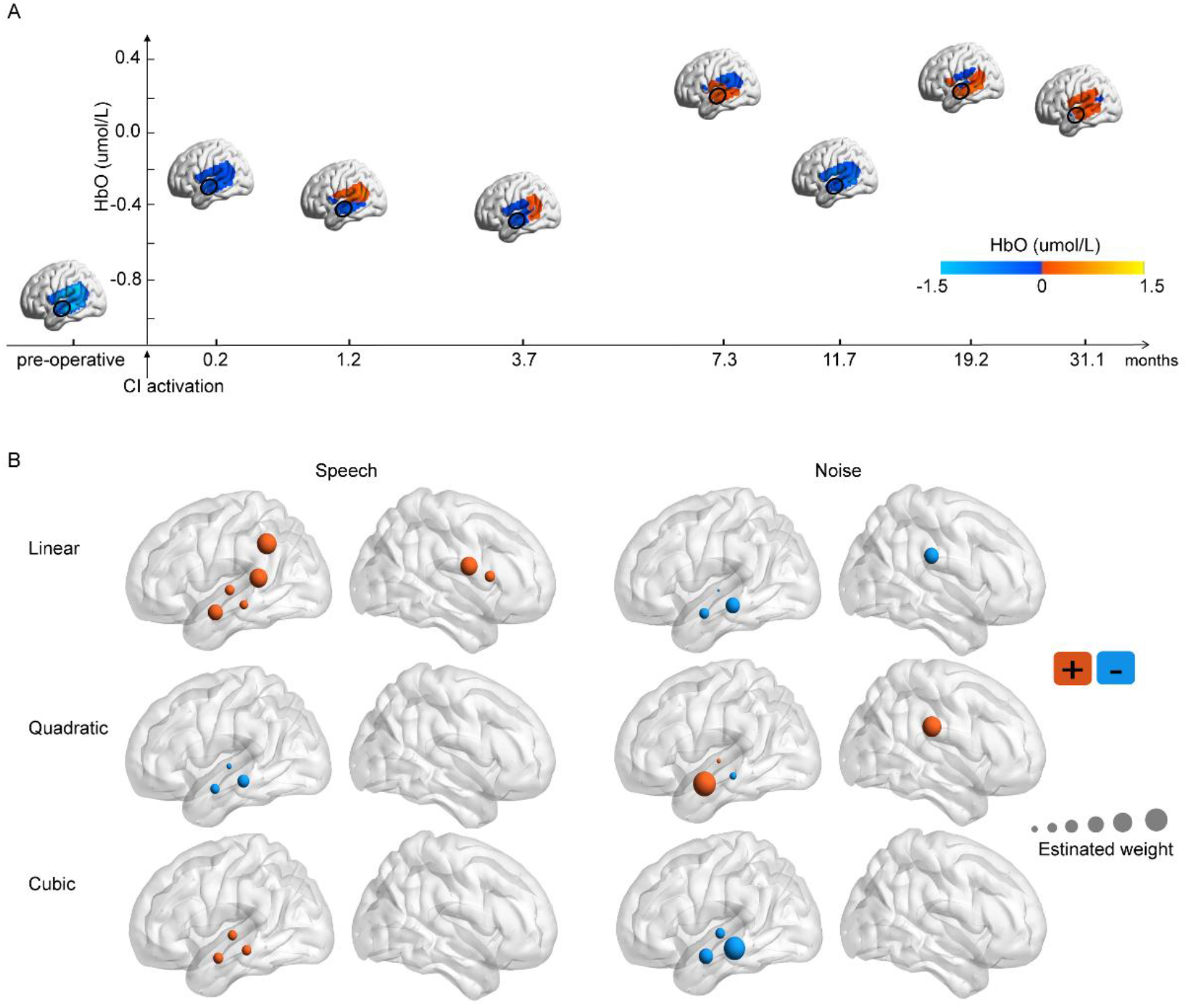
Developmental trajectories of sound evoked responses for cochlear implanted children. (A) Changes of speech evoked HbO responses in the left hemisphere with increasing CI experiences for a sample child participant (age of implantation: 3 years). The y axis marks HbO responses in the left anterior temporal lobe (black circle), which demonstrated a typical non-monotonic, non-linear developmental trajectory. (B) Channel-by-channel results of linear mixed effect (LME) models. A polynomial developmental curve was fitted against the duration of CI experiences up to the third order for each channel and each sound condition. The significant linear (first row), quadratic (second row), and cubic terms (third row) of best fitted models were plotted, with size reflecting estimated weights and color reflecting sign of effect.

To reduce the probability of false positivity caused by multiple comparisons, statistical analyses were conducted on anatomically selected regions of interests (ROIs) (Figure 1B) most relevant for auditory and language processing (Hickok, Gregory et al. 2007, Tellis and Tellis 2016, Poeppel and Assaneo 2020): anterior temporal lobes (ATLs, in red), Sylvian parieto-temporal areas (Spts, in orange), supramarginal gyri (SMGs, in green), and inferior frontal gyri (IFGs, in blue). LME model was conducted on HbO response of each ROI with duration of CI experiences (CI_dur_), sound condition, channel, hemisphere, CI side, age of implantation, residual hearing at CI side and Non-CI side, duration of hearing aid experiences, and gender as candidate fixed factors. However, clinical covariates (CI side, age of implantation, residual hearing in CI side and Non-CI side, duration of hearing aid experiences, and gender) were removed in all optimal models because they have no significant impacts on the development pattern of functional activity (all related effects: p > 0.05).

For ATLs, developmental pattern varied markedly with sound condition (Figure 4A&B; LME: CI_dur_ by condition interactions: *F*s > 16.12, all p values < 0.001), with prominent inter-hemispheric asymmetry (LME: main effect of hemisphere, *F*_(1,2261.6)_ = 23.55, p < 0.001). The left ATL demonstrated oscillatory (cubic polynomial) developmental trajectories for both speech (LME: linear CI_dur_ effect: p < 0.001; quadratic CI_dur_ effect: p = 0.003; cubic CI_dur_ effect: p = 0.007) and noise (LME: linear CI_dur_ effect: p < 0.001; quadratic CI_dur_ effect: p < 0.001; cubic CI_dur_ effect: p < 0.001) processing. Interestingly, speech and noise processing appeared to develop in opposite phases (Figure 4A; LME: CI_dur_ by condition interactions: *F*s > 13.49, all p values < 0.001; main effect of condition, *F*_(1,1109.1)_ = 4.59, p = 0.032), i.e., speech-evoked response peaks coinciding temporally with noise-evoked response troughs. For speech, hemodynamic responses increased after hearing restoration for approximately 9 months, then decreased during the following 12 months before increasing again. For noise, hemodynamic responses decreased after hearing restoration for approximately 9 months, then increased during the following 12 months before decreasing again. In contrast, the right ATL showed no developmental change, with greater HbO responses to speech than to noise (Figure 4B; LME: effect of condition: p = 0.025, detail model parameters see table S1).

**Figure 4.**
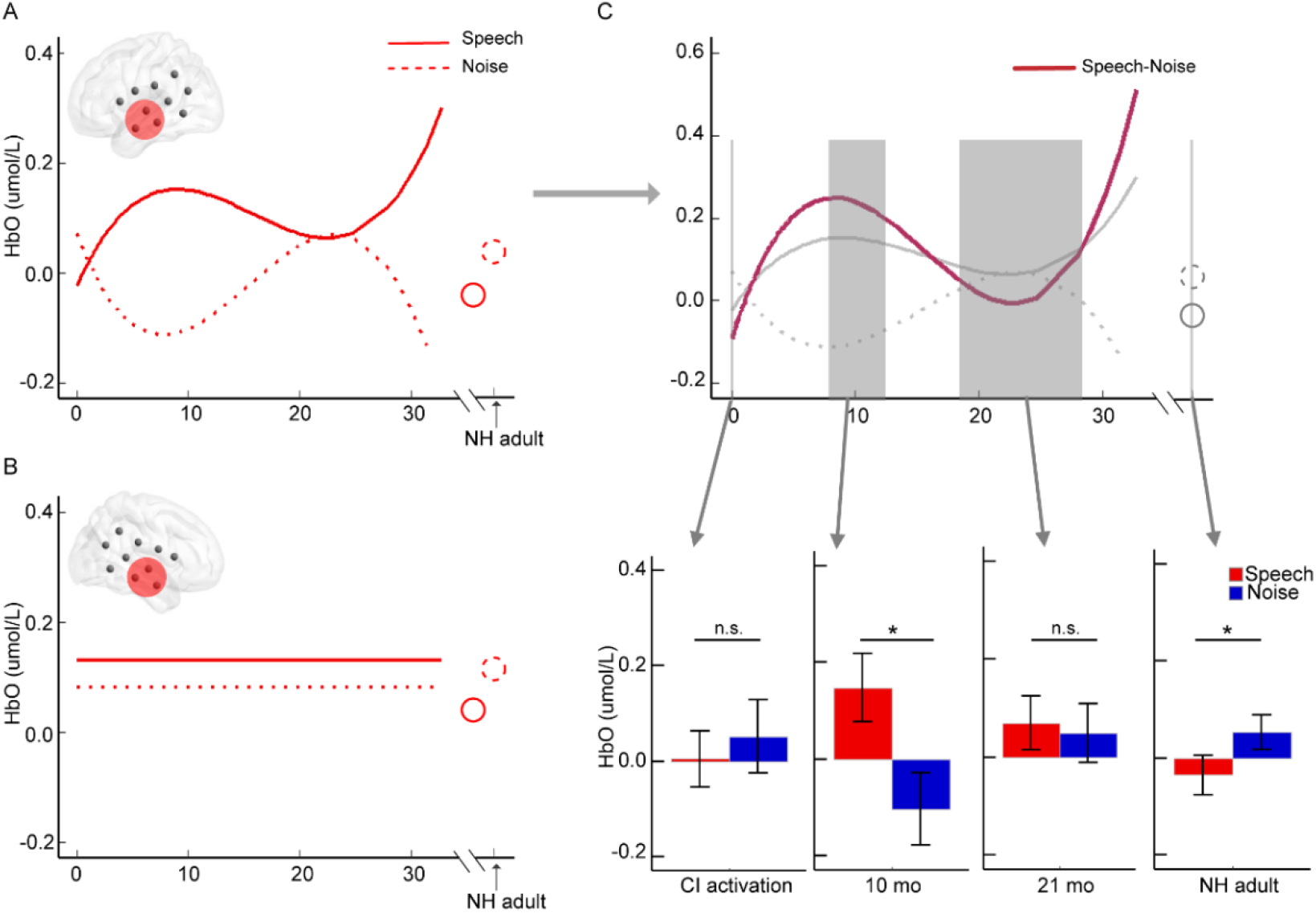
Anti-phasic oscillatory development of speech and noise evoked responses in the left ATL. (A) Fitted developmental trajectory of CI children for speech (solid line) and noise (dotted line) in left ATL (marked in red circle on the inlet brain), illustrating opposite phases for speech and noise development at the group level. As a control, speech (solid circle) and noise (dotted circle) ATL responses from NH adults were shown. (B) Lack of development in right ATL. (C) Developmental coupling of speech and noise responses at the individual level. Upper: Fitted developmental trajectory for within-individual difference in speech and noise evoked responses (marron line) was plotted against the original speech and noise trajectories fitted separately (gray lines), demonstrating that the speech and noise responses changed in opposite phases at the individual level. Bottom: Cortical discrimination of speech (red) and noise (blue) varied across different stages of development (marked in gray bars in the upper panel). Consistent with the oscillatory developmental trajectory, discrimination was not observed at CI activation (N=30) and 21 months after (mean ± std: 21.04±2.57 months, N=16), while was discernable in between with 10 months of CI experiences (mean ± std: 10.09±1.41 months, N=20), similar to NH adults (N=35). Abbreviations: ATL, anterior temporal lobe; CI, cochlear implant; NH, normal hearing. Note: error bars indicate standard errors of mean; n.s., non-significant; *p < 0.05.

To confirm that the anti-phasic development of speech and noise processing occurred at the individual level, LME model was fitted to within-subject difference between speech and noise responses. Consistent with the anti-phasic relationship of the two developmental trajectories at the individual level, another oscillatory developmental trajectory was derived for within-subject speech-noise discrimination (maroon line in Figure 4C and scatterplot see Figure S2; LME: linear CI_dur_ effect: p < 0.001; quadratic CI_dur_ effect: p < 0.001; cubic CI_dur_ effect: p < 0.001), with phases coinciding with the speech and noise trajectories (gray solid and dotted lines in Figure 4C). According to this developmental trajectory, discriminability of speech and noise by cortical responses fluctuated through development, starting from lack of discrimination upon CI activation (Figure 4B; N=30; LME: condition effect: p = 0.531), emergence of discriminability after approximately 10 months of CI experiences with greater speech than noise responses (N=20; condition effect: p = 0.020), temporal reduction of discriminability with further experiences (∼21 months, N=16; condition effect: p = 0.819), and approaching thereafter the discriminability of NH adults achieved by further reduction of speech responses (N=35; condition effect: p = 0.024).

For other ROIs, speech responses increased linearly with CI experiences on the left Spt (Figure 5A; LME: linear CI_dur_ effect: p = 0.013, detail model estimate parameters see table S1) and on the right IFG (Figure 5C; linear CI_dur_ effect: RH, p = 0.013, detail model estimate parameters see table S2), but not in the other regions (all p values > 0.126). Noise responses demonstrated development only on the right SMG, with a U-shaped trajectory (Figure 5B; linear CI_dur_ effect: p = 0.015; quadratic CI_dur_ effect: p = 0.037, detail model estimate parameters see table S2).

**Figure 5.**
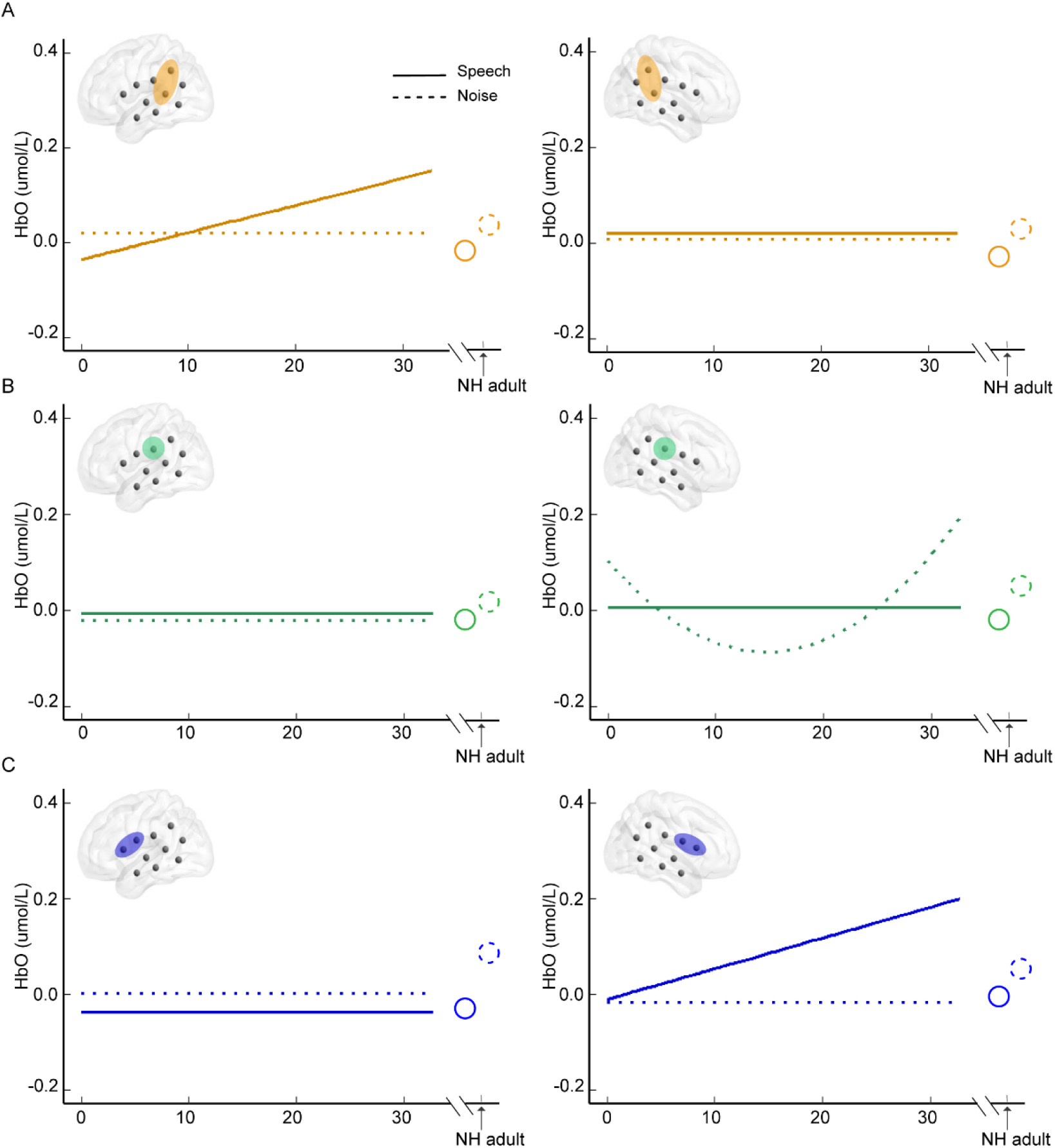
Fitted developmental trajectories in areas other than anterior temporal lobes. (A) Fitted developmental trajectories for speech (solid lines) and noise (dotted lines) in the left (left panel) and right (right panel) Sylvian parieto-temporal areas (marked in orange on the inlet brain illustrations). As a control, speech (solid circle) and noise (dotted circle) responses from NH adults were shown. (B) Fitted developmental trajectories for speech and noise in bilateral supramarginal gyri. (C) Fitted developmental trajectories for speech and noise in bilateral inferior frontal gyri.

For the NH adults, noise evoked greater cortical responses than speech, resulting in speech-noise discrimination in nearly each ROI (Figure S3, all p values < 0.022). In contrast, averaged cortical responses of CI children during the observation period exhibited poorer sound discriminability, achieving significant speech-noise differentiation only in the left ATL (p = 0.018). Note that the left ATL responses of CI children were greater to speech than to noise, reverse to those of the NH adults, indicating that the auditory cortical functions of CI children were far from mature.

### Prediction of CI Outcome by Auditory Cortical Development

Whether the observed cortical developmental changes underlie post-implantation improvement in auditory and communication skills was examined using machine learning (Figure 6A, STAR Methods) in CI children with at least two valid fNIRS assessments and corresponding behavioral assessments (N=47, see table S3 for demographic detail). The CI children showed behavioral improvement in auditory and speech perception skills evaluated by the IT-MAIS/MAIS, CAP, and SIR (Figure 6B; LME: main effect of time: *F*_(2,92)_ = 103.22, p < 0.001). To reduce model overfitting, we performed feature selection procedure on relevant neural data using support vector machine recursive feature elimination (SVM-RFE) algorithm (Martino, Valente et al. 2008) and model evaluation on support vector machine (SVM) classifier using 10-fold cross-validation with 10,000 iterations (Figure 6A). Separate SVM models were built with only clinical features (including gender, age of implantation, CI side, residual hearing in CI side and Non-CI side, duration of CI experiences, and duration of hearing aid experiences, STAR Methods), with only change of HbO responses, and with clinical features in combination with HbO changes. As a control, these models were compared with chance level performance generated by 10000-iteration permutation.

**Figure 6.**
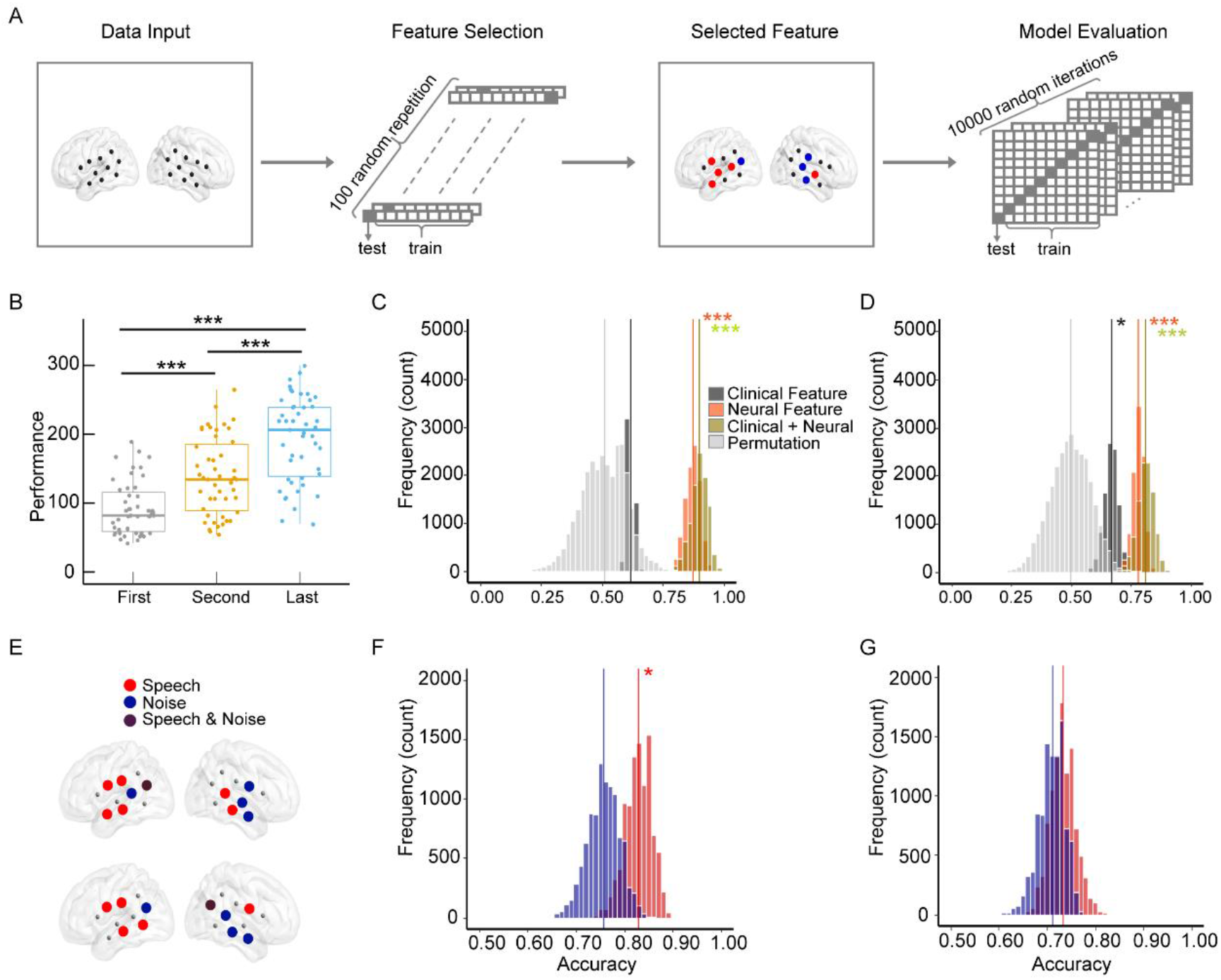
Prediction of post-implantation behavioral improvement of CI children by cortical development. (A) Machine learning procedure for behavioral improvement prediction. CI children were divided into better and poorer groups according to the median development score of questionnaires with guardians. Clinical and/or neural features were entered into a support vector machine (SVM) for group classification. Optimal features were selected using the recursive feature elimination algorithm. Model classification performance was evaluated using ten-fold cross-validation on all data with a bootstrapping test of 10,000 iterations. (B) Behavioral improvement of CI children. As the CI children were not yet able to speak, post-implantation performance on auditory and speech skills were evaluated using summed scores of three questionnaires with guardians. Improvement was observed from approximately one week after CI activation (First, mean ± std: 0.26 ± 0.29 months) to over 2 months of CI wearing (Second, mean ± std: 2.32 ± 3.35 months) and again from then to one year of CI experiences (Last, mean ± std: 12.11 ± 9.62 months). (C) Prediction of behavioral improvement during the first two months. Histograms of classification accuracy from 10,000 iterations were plotted for each of four models. Model with only clinical features (dark gray; input features included gender, age of implantation, CI side, residual hearing in CI side and Non-CI side, duration of CI experiences, and duration of hearing aid experiences) did not outperform chance level (permutation, in light gray), while model with only neural features (orange; changes in speech and noise responses in all fNIRS channels as input) did so by far, with similar performance to model of mixing clinical and neural features (yellow green). (D) Prediction of behavioral improvement during the first twelve months. (E) The selected neural features in optimal models for prediction of improvement in the first 2 months (top row) and the first 12 months (bottom row). (F) Prediction of behavioral improvement during the first 2 months using only speech (red) or noise (blue) evoked responses. (G) Prediction of behavioral improvement during the first 12 months using only speech (red) or noise (blue) evoked responses. Note: * p < 0.05, *** p < 0.001.

During the first 2 months of CI experiences (from the mean of 0.26 months to 2.32 months after CI activation; Figure 6C), prediction accuracy of behavioral improvement by clinical features was 0.62 (black histogram), not different from chance level (Z test: *Z* = 1.20, p = 0.230). HbO changes yielded classification accuracy of 0.87 in isolation and 0.90 in combination with clinical features, significantly higher than clinical features (Z test: *Z* > 19.51, all p values < 0.001). Throughout the observation period (from the mean of 0.26 months to 12.11 months after CI activation, Figure 6D), clinical features yielded prediction accuracy of 0.67, slightly higher than chance level (black histogram, *Z* = 2.01, p = 0.044). Prediction accuracy of HbO changes was 0.78 in isolation and 0.81 in combination with clinical features (Figure 6D, orange and green histograms), also significantly higher than predictability of clinical features (*Z* > 3.91, all p values < 0.001).

To elucidate the relative predictability of speech and noise evoked responses, their changes were entered separately into the SVM classifier. During the first 2 months of CI experiences (Figure 6F), the classification accuracy was 0.83 for speech and 0.76 for noise responses (Z test: *Z* = 2.16, p = 0.031), with selected channels mostly on the left hemisphere for speech and on the right hemisphere for noise condition (Figure 6E, top row). Throughout the observation period (Figure 6G), the classification accuracy was 0.73 for speech and 0.71 for noise responses (Z test: *Z* = 0.88, p = 0.379), with slightly altered channel selection (Figure 6E, bottom row). Overall, for speech, classification accuracy of early change of cortical responses was higher than the entire period (speech: *Z* = 3.47, p < 0.001; noise: *Z* = 1.42, p = 0.156), indicating early neural change may be more sensitive to change in behavioral performance.

## DISCUSSION

The current study delineated early development of auditory cortical functions for young children with prelingual bilateral hearing loss from the onset of hearing with cochlear implant (CI). At CI activation, CI children showed general lack of sound responses except in the right anterior temporal lobe and lack of speech-noise discrimination, functionally inferior to the healthy brain even before birth. Longitudinal fNIRS recordings of the CI children up to 32 months, a critical period before speech acquisition, revealed stimulus specific developmental patterns across the auditory and speech processing regions. Most remarkably, cortical responses in the left anterior temporal lobe (lATL) changed with increasing hearing experiences in an oscillatory manner, with speech and noise processing developing in opposite phases. Further, speech responses increased linearly in the left sylvian parieto-temporal area (lSpt) and the right inferior frontal gyrus (rIFG), while noise responses showed a U-shaped developmental pattern in the right supramarginal gyrus (rSMG). Cortical developmental changes were capable of predicting post-implantation improvement on auditory and communication performance using an SVM classifier, with an advantage of speech over noise processing during the first 2 months of hearing experiences that disappeared in about one year. Overall, the results provide the first longitudinal evidence of how the very early hearing experiences shape auditory and speech functions of the human brain before speech acquisition.

### Effect of Auditory Deprivation Early in Life

In the current study, young CI children with prelingual bilateral deafness were used as a model of early auditory development before speech acquisition. Suggesting that their brain was in a very early stage of auditory functional development, at onset of CI hearing, their auditory and speech processing areas were irresponsive to sounds except in the right ATL, and were incapable of discriminating sound types (speech vs. noise). In contrast, the healthy brain at birth was able to recognize their mother’s voice from unfamiliar voices (Decasper and Fifer 1980, Moon, Christine et al. 2000), native from nonnative vowels (Moon, C. et al. 2013), and different consonants (Cabrera and Gervain 2020). Even before term birth, left superior temporal and supramarginal regions of preterm infants (25-37 weeks of post menstrual age) are already activated by speech sounds (Baldoli, Scola et al. 2015), and inferior frontal regions of preterm infants (28-32 weeks of gestational age) are able to detect change of human voice and change of phoneme (Mahmoudzadeh, Dehaene-Lambertz et al. 2013). Indeed, the human brain started to respond to auditory stimuli at 24 weeks of gestation in utero (Birnholz and Benacerraf 1983). Notably, increasing evidence in in-vivo fetal neuroimaging indicates that auditory development is affected by sensory input before birth and primordial language networks develop throughout the gestational stage (for a recent review, Ghio, Cara et al. 2021). Thus, the current results of CI children indicate that auditory deprivation early in life delays the development of auditory cortical functions, resulting in prominent inferiority to the normal-hearing brain even at birth.

The cortical response patterns of CI children were consistent with animal models of congenitally deaf cats (Kral and Sharma 2012, Kral and A. 2013, Kral, Dorman et al. 2019). Compared with hearing cats, deaf cats showed delayed postnatal synaptogenesis, with little evoked synaptic activity in the auditory cortex at birth. Upon hearing restoration with CI, the auditory cortex exhibited a “catch up” stage of accelerated synaptic growth and exaggerated pruning, with peak neural activity surpassing the hearing cats (overshooting). Despite of the delay caused by auditory deprivation, the developmental trajectories of auditory cortical activity are of similar shape between hearing and CI cats. As many of the CI children would ultimately acquire speech (Anderson, Wiggins et al. 2017, Feng, Ingvalson et al. 2018), it is highly possible that their brain might undergo similar developmental catch up with CI hearing experiences, making them an ideal model to study early auditory development of the human brain preceding speech acquisition.

Interestingly, after auditory deprivation early in life, the only cortical region responsive to sounds is the right ATL (Figure 2C & S1). This observation is consistent with previous brain imaging demonstrations that the normal hearing neonates showed clear right hemisphere dominance for non-linguistic aspects of auditory processing (Perani, Saccuman et al. 2010, Arimitsu, Uchida-Ota et al. 2011, Bisiacchi and Cainelli 2021, Bourke and Todd 2021). In normal hearing adults, the rATL is associated with voice and speaker identity processing (Kriegstein and Giraud 2004, Elia, Formisano et al. 2008, Andics, Mcqueen et al. 2010, Bestelmeyer, Belin et al. 2011, Andics, Mcqueen et al. 2013, Bonte, Hausfeld et al. 2014). The preservation of the functionality of rATL with auditory deprivation thus possibly enables deaf children to recognize familiar voice immediately upon hearing restoration, a skill exhibited by the normal hearing brain at birth (Mahmoudzadeh, Dehaene-Lambertz et al. 2013). The rATL responses were not accompanied by responses in the auditory cortex on either hemisphere. This could result from methodological limitation, as considerable portion of the auditory cortex lies inside the Heschl’s gyrus (HG) and probably escapes detection of fNIRS (Cui, Bray et al. 2011).

### Development of Auditory Cortical Functions with Restored Hearing Experiences

With restored hearing experiences (up to 32 months), auditory cortical responses of CI children showed differential developmental patterns for speech and noise. For speech, developmental changes were observed in lATL, lSpt, and rIFG (see Figure 4 & Figure 5), while for noise, development occurred in lATL and rSMG (see Figure 4 & Figure 5).

The speech developing areas on the left hemisphere, lATL and lSpt, lie respectively in the ventral and dorsal streams in terms of the celebrated dual-stream theory of auditory and speech processing (Hickok and Poeppel 2007, Rauschecker and Scott 2009, Poeppel and Assaneo 2020). According to this theory, the ventral stream is responsible for sound recognition and speech comprehension, while the dorsal stream is responsible for auditory-motor integration. Relatively little is known about the functional development of the two processing streams early in life. The current study with CI children indicates that both streams started to develop from hearing onset, suggesting that the brain was striving to acquire “listening” and “speaking” skills simultaneously. Development was only observed in the temporal lobe and neighboring parietal areas, not in the frontal speech areas, consistent with the “temporal-to-frontal” developing order suggested by DTI studies (Perani, Saccuman et al. 2011, Brauer, Anwander et al. 2013) and by EEG/MEG studies with young children (Skeide and Friederici 2016). The rATL was not considered part of the dual-stream language network (Hickok and Poeppel 2007, Rauschecker and Scott 2009, Poeppel and Assaneo 2020). For NH adults, fMRI studies revealed that rIFG was involved in multiple functions ranging from prosodic processing of sounds (Meyer, Kai et al. 2002, Friederici 2011, Skeide and Friederici 2016) to executive processes (Hampshire, Chamberlain et al. 2010, Madsen, Baare et al. 2010, Banich and Depue 2015, Gavazzi, Giovannelli et al. 2021). The early development of rIFG in CI children likely reflects emergence of prosodic processing, a skill that is critical for CI hearing (Velde, Schiller et al. 2019) and language acquisition(Gerken 1996, Fló, Brusini et al. 2019) and that start to develop within the first year of life for NH children (Homae, Watanabe et al. 2007, Jusczyk, Cutler et al. 2010).

In terms of noise processing, the development of lATL is expected, as the area is regarded an important stage in the ventral stream responsible for sound recognition (the “what” pathway). Evidently, the early development of lATL for both speech and noise would lead to discrimination of sound types. The rSMG, in the NH adult brain, is involved in affective aspects of sensory processing, including emotion and empathy (Singer, Seymour et al. 2004, Lamm, Decety et al. 2011, Belyk, Michel et al. 2014, Hoffmann, Koehne et al. 2016, Esménio, Soares et al. 2019, Wada, Honma et al. 2021). This appears to be the case in neonates, where the rSMG showed heighted fNIRS sensitivity to emotional than neutral prosodies (Zhang, Zhou et al. 2017, Zhang, Chen et al. 2019). Thus, the rSMG and rIFG appear to change in synergy, allowing the right hemisphere to capture non-linguistic, specifically affective and prosodic, information during early stage of auditory development.

Overall, the current study indicates a clear distinction in developmental emphases for the left and right hemisphere from the onset of (restored) hearing: The left hemisphere, primarily the areas next from the auditory cortex in the ventral and dorsal streams of speech processing, appears to strive to acquire listening and speaking abilities concurrently, while the right hemisphere appears responsible for non-linguistic aspects of auditory processing. Thus, the current results provide longitudinal developmental evidence for theories of functional asymmetries of the brain (Perani, Saccuman et al. 2010, Arimitsu, Uchida-Ota et al. 2011, Bisiacchi and Cainelli 2021, Bourke and Todd 2021, Hartwigsen, Bengio et al. 2021).

### The Anti-phasic Oscillatory Development of Speech and Noise Processing

Oscillatory or cyclic developmental trajectories of brain and cognitive functions, among many other possibilities, are theoretically expected but rarely observed, due to the scarcity of longitudinal data that can separate individual and group level variances (Sanchez-Alonso and Aslin 2020). The closest example that we are aware of is from resting state EEG studies with hundreds of children (Thatcher, Walker et al. 1987, Thatcher 1992, Thatcher, North et al. 2008). Phase and coherence of EEG signals across different scalp-placed electrodes, computed as an index of resting network connectivity, revealed multiple cycles of changes from infancy to childhood and adolescence (Thatcher, Walker et al. 1987, Thatcher 1992, Thatcher, North et al. 2008). However, rapidly accumulating data acquired with non-invasive brain imaging during the past decades, particularly structural MRI, typically reveal bell (or U) shaped trajectories for brain development. For example, gray matter density in most of the neocortex increased during early childhood and then decreased until adulthood (Sowell, Peterson et al. 2003, Sowell, Thompson et al. 2004, Shaw, Kabani et al. 2008, Raznahan, Lerch et al. 2011, Raznahan, Shaw et al. 2011, Coupe, Catheline et al. 2017), with peaking age varying across brain regions and linked to cognitive functions (Shaw, Greenstein et al. 2006) or disorders (Alexander-Bloch, Reiss et al. 2014). The turning point of the bell-shaped trajectories is still under debate, as many recent large-scale MRI studies revealed monotonic decrease of cortical thickness after 3 to 4 years old, placing the peaking age to infancy and toddlerhood (for a recent review, Walhovd, Fjell et al. 2017). Similarly, diffusion tensor imaging (DTI) data have delineated U-(radial diffusivity) or inverted U-shaped (fractional anisotropy) curves in major white matter tracts, with volume peaking from late adolescence to middle life (Lebel and Beaulieu 2011, Coupe, Catheline et al. 2017). Developmental trajectories from functional imaging are rarely available, particularly through the first years of life (Madhyastha, Peverill et al. 2017, Lana, Abaci et al. 2018). As structural-functional coupling of the developing brain remains largely unknown and likely changing through life (Passingham, Stephan et al. 2002, Bennett and Rypma 2013, Vasung, Abaci Turk et al. 2019, Wang, Ghaderi et al. 2021), functional developing trajectories cannot be directly derived from structural trajectories.

The current study provides a first example of oscillatory functional development with longitudinal brain imaging data, echoing the previous observation with EEG coherence (Thatcher, Walker et al. 1987, Thatcher 1992, Thatcher, North et al. 2008). Given that synaptic connections are considered the major neuronal events associated with sensory evoked hemodynamic responses (Vasung, Abaci Turk et al. 2019), we suggest that the oscillatory trajectories reflect alternating dominance of progressive (i.e., synaptic growth) and regressive (i.e., experience-driven pruning) processes of synaptic connectivity, consistent with the proposed mechanism for cyclic EEG trajectories (Thatcher, North et al. 2008). Across the life span, synaptic connections in the primate brain also exhibit a bell-shaped developmental curve, with exuberant growth during the first years of life to peak at nearly twice as many as in the adult brain before declining gradually throughout childhood and adolescence (Casey, Tottenham et al. 2005, Stiles and Jernigan 2010). Recent advances in real-time imaging indicate that formation and elimination of synaptic connections take place concurrently in the developing brain at the microscopic time scale of minutes to hours, subject to modulation of neural activity (Hua and Smith 2004, Faust, Gunner et al. 2021). In terms of developmental time scale, the current oscillatory trajectories turned from increasing to decreasing or from decreasing to increasing at the rate of approximately 10 to 15 months, falling somehow in between the previously documented macroscopic (years to lifespan) and microscopic (minutes to hours) development of synaptic connections. In terms of brain location, oscillatory trajectories for sound evoked hemodynamic responses were observed only in the left ATL of young CI children, a region critical for sound recognition (Hickok and Poeppel 2007) and a network connection hub for high-level cognitive functions (Ralph, Jefferies et al. 2016) in the mature brain. Taken together, the current data suggest that the macroscopic brain development may consist of mesoscopic alternations of synaptic growth and pruning, particularly during the first years of life, as a regional mechanism to boost experience-driven formation of neural circuits for critical functions.

Most interestingly, the left ATL responses changed with increasing hearing experiences in an anti-phasic fashion for speech and noise. The anti-phasic relationship was observed at the individual level (Figure 4), revealing a novel form of developmental coupling of different functions in the same brain region. Such coupling could reflect synergic formation of different subsets of neural circuits for speech and noise, i.e., growth in one set being synchronized with pruning of the other set. Alternatively, it could be a manifestation of stimulus specific computations carried out by the same circuits, such as increased activity for one sound and decreased activity for the other. The developmental phase differences led to fluctuations of cortical discriminability of the two sound types with time of measuring, opposing the widely held assumption of monotonic improvement of functional discrimination during development. This phenomenon may help reconcile conflicting results obtained via cross-sectional comparisons, such as cortical discrimination of speech and backward speech in the left temporal lobe of infants at 3 months of age (Dehaene-Lambertz, Dehaene et al. 2002) but not at 4 months (Sato, Sogabe et al. 2010). Despite of such fluctuations, developmental coupling of different circuits or their computations is of evident ecological advantage, allowing for simultaneous and coordinated emergence/maturation of multiple functions under limited experiences.

## STAR METHODS

### KEY RESOURCES TABLE

| REAGENT or RESOURCE | SOURCE | IDENTIFIED |
| --- | --- | --- |
| **Software & Algorithms** |  |  |
| MATLAB | Mathworks | https://www.mathworks.com/ |
| R | R Core Team, 2019 | https://www.r-project.org/ |
| python | Python | https://www.python.org/ |
| Praat | David 2008 | http://www.praat.org/ |
| FC_NIRS | Xu, Liu et al. 2015 | https://www.nitrc.org/frs/?group_id=883 |
| HOMER2 | Huppert et al. 2009 | https://www.nitrc.org/frs/?group_id=619 |
| BrainNet Viewer | Xia et al.2013 | https://www.nitrc.org/projects/bnv/ |
| Custom code for this paper | N/A | https://github.com/wumeiyunwmy6/CI_brain-development |
| Deposited Data | N/A | https://osf.io/ngkv2/ |

### RESOURCE AVAILABILITY

#### Lead contact

Any additional information and requests for resources or data should be directed to and will be fulfilled by the Lead Contact, Yu-Xuan Zhang at.

#### Materials availability

This study did not generate new unique materials.

#### Data and code availability

Raw data that support the conclusions of this study are available for public download in https://osf.io/ngkv2/, as well as analyzed data and codes are available in Github at https://github.com/wumeiyunwmy6/CI-development-cortical-response.

### EXPERIMENTAL MODEL AND SUBJECT DETAILS

#### Participants

Children with severe-to-profound prelingual bilateral sensorineural hearing loss were recruited from the Peking University First Hospital before cochlear implantation surgery (CI children, N=135; mean + std age: 3.02 + 2.17 years, 58 females). Normal hearing (tone threshold ≤ 20 dB HL across 0.25–16 kHz at both ears) young adults (NH adults, N=35; mean + std age: 22.76 + 1.78 years, 20 females) were recruited as a control of the experimental protocol. Informed consent was obtained from parents (or guardian) of CI children and NH young adults before the experiment. This study was approved by the Ethical Committee of Peking University and the Peking University First Hospital (protocol number: research #239).

All child participants complied with the cochlear implantation criteria set out by China National Guidelines for Cochlear Implantation (Committee 2014). Their electrodes were estimated by the clinical team using neural response telemetry (NRT) during the operation. Out of the 135 child participants, 51 were unable to finish at least one session of fNIRS testing with valid data. The current study included only children with either valid evaluation at CI activation or at least two repeated measures thereafter (224 fNIRS measurements from 67 child participants). The distribution of individual longitudinal track for each child was shown in Figure S1A. The fNIRS data preprocessing and processing procedure was shown in Figure S4 and the demographic information of the CI children included in the current analyses was in table S3 (characteristics of each CI child see: https://github.com/wumeiyunwmy6/CI-development-cortical-response).

### METHOD DETAILS

#### Experimental design

Auditory cortical development of CI children was examined in a longitudinal developmental design using repeated functional near-infrared spectroscopy (fNIRS). On each hospital visit after implantation, cortical responses to different sound conditions were recorded using fNIRS in the auditory neuroscience laboratory at Beijing Normal University, and auditory and speech skills were evaluated using questionnaires with guardians. A recommended schedule for laboratory tests was provided according to the standard clinical protocol. Actual test frequency and timing were decided by the guardians. Given the uncertainty in developmental stage of the CI children, only NH young adults were recruited to provide a control of the matured brain using the same imaging protocol.

#### CI outcome assessment

Most CI child participants were toddlers without effective communication abilities. Consistent with current clinical practice, CI outcome on auditory and speech skills was measured using the Infant-Toddler Meaningful Auditory Integration Scale (IT-MAIS/MAIS) (Zimmerman-Phillips 1997), Categories of Auditory Performance (CAP) (Archbold, Lutman et al. 1995), and Speech Intelligibility Rate (SIR) (Allen, Nikolopoulos et al. 1998), which are suitable to the young children, from the children’s parent/ guardian. The normalized sum of these questionnaires was calculated to provide a unified measure for the auditory and speech ability of children.

#### fNIRS recording

At each laboratory visit, the child participant was seated on a child chair or the lap of their guardian in a sound-attenuated booth, watching a muted cartoon video or playing with silent toys, while an elastic fNIRS cap (head size 52 cm, EASYCAP GmbH, Woerthsee-Etterschlag, Germany) was placed on their head. The cap was fitted with 8 light sources and 8 light detectors (3 cm between neighboring sources and detectors), generating 20 measurement channels covering the temporo-frontal auditory and language related areas on both hemispheres (Hickok, Gregory et al. 2007, Tellis and Tellis 2016, Poeppel and Assaneo 2020) (Figure 1B). Probe placement was in accordance with the international 10–10 system.

Each fNIRS session consisted of 5 minutes of resting state followed by approximately 17 minutes of passive listening. Imaging data were recorded using NIRSport2 optical topography systems (NIRx Medical Technologies, NY, USA) at wavelengths of 760 nm and 850 nm, with a sampling rate of 7.81 Hz. Cochlear implants and hearing aids if in wearing were turned off for the resting state and on for passive listening. Sounds were presented using custom programs with Psychtoolbox for Matlab (Brainard 1997, Kleiner, Brainard et al. 2007) from two cube BOSE loudspeakers (BOSE, USA) speakers placed at 45º left and right to the direction the participant was facing (Figure 1C). The sound level at the participant’s head position was 65 dB SPL.

Passive listening consisted of 20 blocks of sound presentation from 4 conditions (5 blocks/condition) in a pseudo-randomized order, excluding consecutive blocks from the same condition. Each block lasted for 20 seconds, followed by a silence interval of 20 to 25 seconds. To relieve the child participant’s physical or mental tiredness, the 20 blocks were divided into three sessions, allowing the child to rest between sessions. The experiment would stop whenever the participant showed crying, emotional disturbance, frequent head movements, or at the request of the guardian. The NH adults were recorded using the same protocol as the CI children, except for an adult-size cap (58 cm).

fNIRS were recorded for four sound conditions: natural speech, instrumental music, multi-talker babble noise, and speech-in-noise (for stimulus samples see https://github.com/wumeiyunwmy6/CI-development-cortical-response). Only the speech and noise conditions were analyzed in the current study (for sound waveforms and amplitude envelopes, see Figure 1A). The speech stimuli were five 20-second segments adapted from child audio books using Praat software (David 2008). Noise was prepared by mixing audio tracks of 6 stories by different tellers (for detailed procedure see Gao, Yan et al. 2020).

### QUANTIFICATION AND STATISTICAL ANALYSIS

#### Transcranial localization of fNIRS channels

To improve the accuracy of channel locations, we took advance of a recent advance in transcranial brain atlas for children (Zong, Zheng et al. 2021) to assess cortical locations of the fNIRS channels *post hoc*, using a child head manikin (for detailed procedure, see (Zhao, Xiao et al. 2020). First, five anatomical landmarks (NZ: Nasion, IZ: Inion, CZ: Vertex, AL/AR: left/right preaurical point) and twenty-five sparse sampling points of scalp surface were recorded using a 3D digitizer to construct the Continuous Proportional Coordinate (CPC) system. Second, all fNIRS channels were projected onto the CPC system based on the scalp landmark to determine their CPC coordinates. The CPC coordinates were then transformed to MNI coordinates and BA labels (see Table S4 for detail) using the transcranial brain atlas (TBA) of young children (Zong, Zheng et al. 2021). In the current study, cortical responses were mapped to the brain according to the child TBA-based localization, as TBA-based method results in a more optimized probe arrangement compared to the traditional 10/20-based method (Zhao, Xiao et al. 2020).

#### fNIRS data preprocessing

There are typically abundant head motion artefacts in fNIRS recordings of toddlers (Smith, Condy et al. 2020). We first performed data quality screening (Figure S4), using FC_NIRS (Xu, Liu et al. 2015) and custom scripts in Matlab (The MathWorks, USA). Recordings of a given channel were regarded as invalid and excluded from further analysis if heartbeats (1∼1.5 Hz) were not detectable or the coefficient of variation exceeded 20% (Piper, Krueger et al. 2014). A given fNIRS recording was regarded as valid if no more than 30% channels were excluded (Bulgarelli, Klerk et al. 2020), and were then preprocessed using the HOMER2 toolbox (Huppert, Diamond et al. 2009). Motion artifact correction was conducted using combination of wavelet and spline methods recommended for children (Lorenzo, Pirazzoli et al. 2019). A band-pass filter of 0.01 to 0.2 Hz was applied to minimize physiological noise such as breathing and improve signal-to-noise ratios. The relative concentration changes of oxy-hemoglobin (HbO) and deoxy-hemoglobin (Hb) were calculated by the Modified Beer-Lambert Law with differential path length factor set at 6 for the two wavelengths and the 5-second silence preceding stimulus onset for baseline correction.

#### Cortical hemodynamic response quantification

In the current study, cortical hemodynamic responses were quantified using the oxy-hemoglobin signal, as it is known to be more strongly correlated with cognitive activity (Malonek, Dirnagl et al. 1997, Strangman, Culver et al. 2002, Fu, Mondloch et al. 2014) and more sensitive to the regional cerebral blood flow changes (Hoshi 2007, Fu, Mondloch et al. 2014). HbO response for each stimulus block was calculated as averaged HbO concentration from 5 to 20 seconds after sound onset, which was then averaged across blocks for each stimulus condition. Averaged HbO concentration was regarded more suitable to children than beta coefficients calculated based on the adult hemodynamic response function (Shultz, Vouloumanos et al. 2014, Issard and Gervain 2018, Smith, Condy et al. 2020).

#### Statistical analyses

Outlying HbO responses, i.e., more than three standard deviations away from mean of each condition, were excluded. Statistical analysis was performed in R environment (Team 2015), including t test and linear mixed effect (LME) models. Specifically, ttest function was used to compare cortical activation of sounds versus silence, as well as to compare CI children and NH adults. False Discovery Rate (FDR) was used in multiple comparisons to correct the P-value using the fdrtool package (Klaus and Strimmer 2015).

Developmental trajectories were estimated using LME models with the lmer function in the lmerTest package (Kuznetsova, Brockhoff et al. 2016), fitted by maximum likelihood and P values calculated by t tests with Satterthwaite’s method. For each cortical region, a full LME model was built, with cortical hemodynamic response as the dependent variable, duration of CI experiences (CI_dur_) as the continuous predictor, condition, channel, hemisphere as fixed factors. Subject-level variations in intercept and CI_dur_ effect were included, to separate group and individual effects (Smith, Condy et al. 2020). Interaction items without significant effects would be removed from the model. For each developmental trajectory, up to the cubic term of polynomial functions on CI_dur_ were fitted and best fitted models were selected using Akaike information criterion (AIC) and log-likelihood (logLik). Clinical and demographic factors, including CI side, age of implantation, residual hearing in CI side and Non-CI side, duration of hearing aid experiences and gender, were entered into the optimal models as covariates to test their influence (see table S5 and S6 for all model contrast and table S7 for optimal mixed-effects model).

#### Neuro-behavior relation analyses

The support-vector-machine (SVM) classifier in the python environment was used to explore the predictability of clinical features and cortical responses for CI outcome during two periods: the first 2 months and the first year of CI experiences. Clinical features included duration of CI experiences, CI side, age of implantation, residual hearing in CI side and Non-CI side, duration of hearing aid experiences, and gender. Neural features included changes in speech evoked and noise evoked HbO responses at two tests in all channels. Missing HbO values were filled in using averaged responses from neighboring channels. Clinical and neural data were converted into normalized S×V matrices, where S is the number of children and V is the number of features. CI outcome was calculated as changes in summed questionnaire scores at two tests divided by time lapse. CI children were divided into better (N=24) and poorer (N=23) groups using the median CI outcome.

All participants were randomly divided into training (90%) and testing (10%) datasets. Features with highest classification accuracy were selected using SVM-RFE algorithm with ten-fold cross-validation. Model performance with the optimal feature subset was evaluated using the SVM classifier with linear kernel and the default parameters. The area under the ROC curve (AUC), sensitivity, accuracy, and precision were calculated from classification performance of the models. Mean performance of ten-fold cross-validation on all data with a bootstrapping test of 10,000 times was calculated using the scikit-learn package (Pedregosa, Varoquaux et al. 2012) combined with custom scripts. A permutation test with 100 repetitions was conducted to compare classification accuracy with chance, generating Z-score statistics for condition difference.

#### Data Visualization

Brain surfaces were visualized using the BrainNet (Xia, Wang et al. 2013).

## SUPPLEMENTAL INFORMATION

## ACKNOWLEDGEMENTS

We thank all CI children and their parents/guardians who participated in our experiment. We thank Yi Yu for help with drawing graphic abstracts. This work was supported by the National Natural Science Foundation of China (82071070), Beijing Municipal Science & Technology Commission (Z191100006619027), and Beijing Brain Initiative of Beijing Municipal Science & Technology Commission (Z181100001518003).

## AUTHOR CONTRIBUTIONS

Y.Z. and Y.L. conceived and designed research; M.W. Y.W, W.W, H.L, T.X, S.W, M.L., C.W. collected data; M.W. and X.Z. and Y.W. analyzed data; C.Z. methodology; Y.Z. and M.W. interpreted the results and wrote the paper.

## Supplementary Information

This file includes:

Supplemental Figures (Figures S1 to S4)

Supplemental Tables (Tables S1 to S7)

## Supplemental Figures

**Figure S1.**
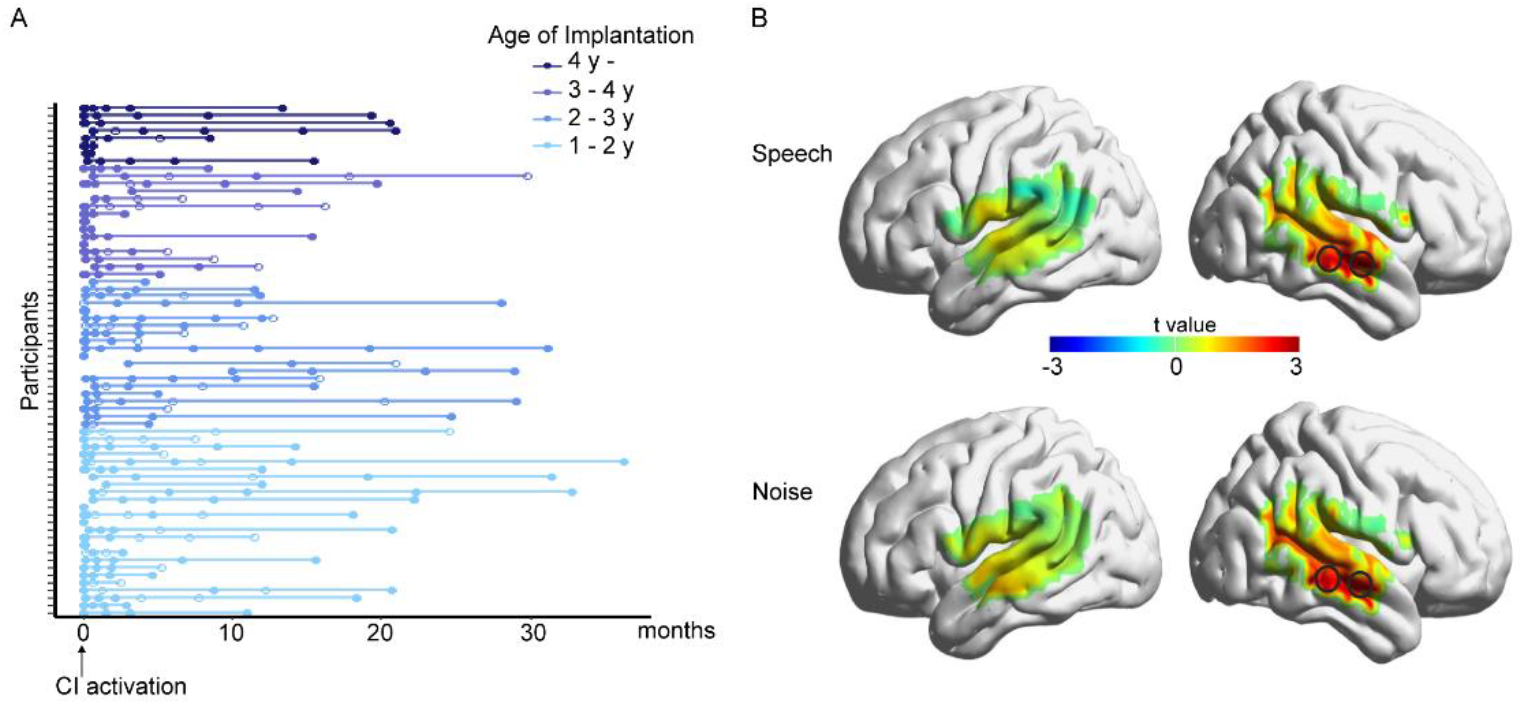
Related to Figure 2. Testing schedule and sound responsiveness at hearing onset of the cochlear implanted (CI) children. (A) Testing schedule of 67 CI children. Each line represents an individual child and each circle represents an fNIRS assessment (filled circles indicate valid data and open circles indicate invalid data). Color indicates range of the age of implantation (y: year). (B) T statistics map of cortical responses elicited by speech (top) and noise (bottom) in comparison with silence at CI activation (N=30). Significant activation (p<0.05 FDR corrected) was marked with black circles.

**Figure S2.**
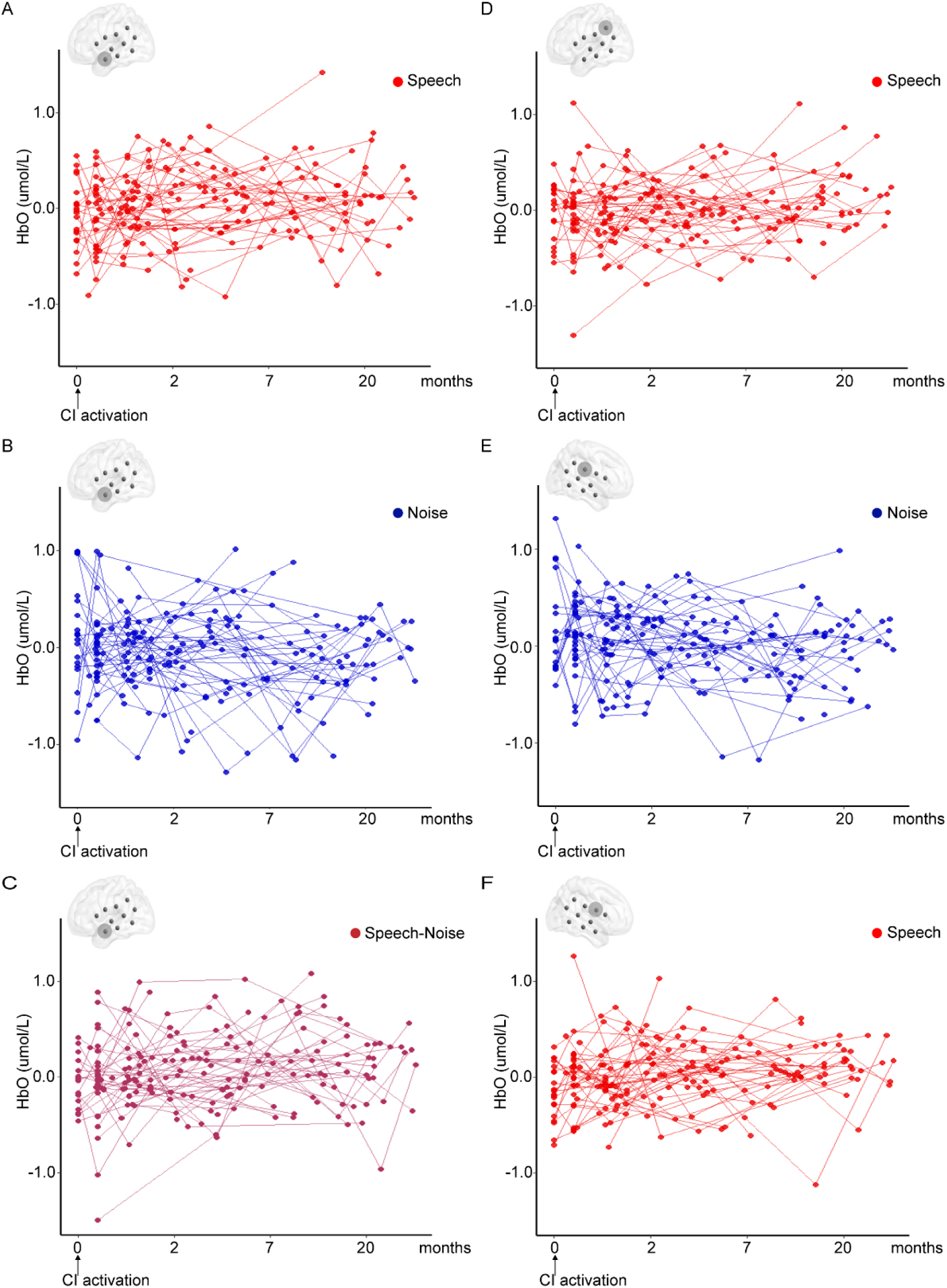
Related to Figure 4 and 5. Sample raw developmental changes of CI children (N=57).

**Figure S3.**
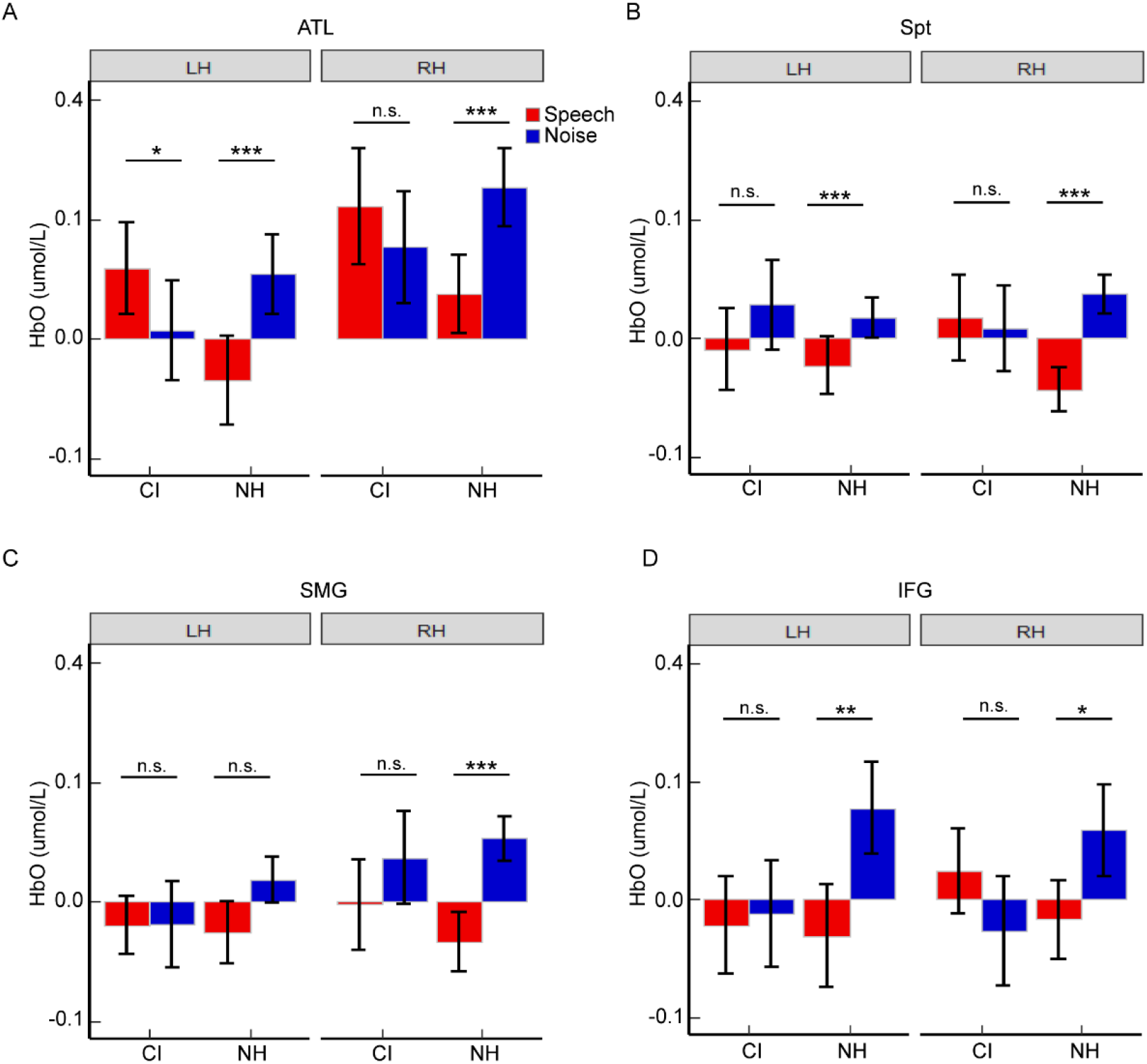
Related to Figure 4 and 5. Cortical speech-noise discrimination of CI children and NH adults in ATL (A), Spt (B), SMG (C), and IFG (D). NH adults showed greater noise than speech responses in all of the regions of interest except the left SMG. In contrast, CI children (averaged across the observation period) showed significant discrimination only in the left ATL, but in opposite direction (greater to speech than noise) to that of NH adults. Abbreviations: CI, Cochlear implanted; NH, normal hearing; ATL, anterior temporal lobe; Spt, sylvian parieto-temporal areas; SMG, supramarginal gyrus; IFG, inferior frontal gyrus; LH=Left hemisphere, RH=Right hemisphere. Note: n.s., non-significant, *p < 0.05, **p < 0.01, ***p < 0.001.

**Figure S4.**
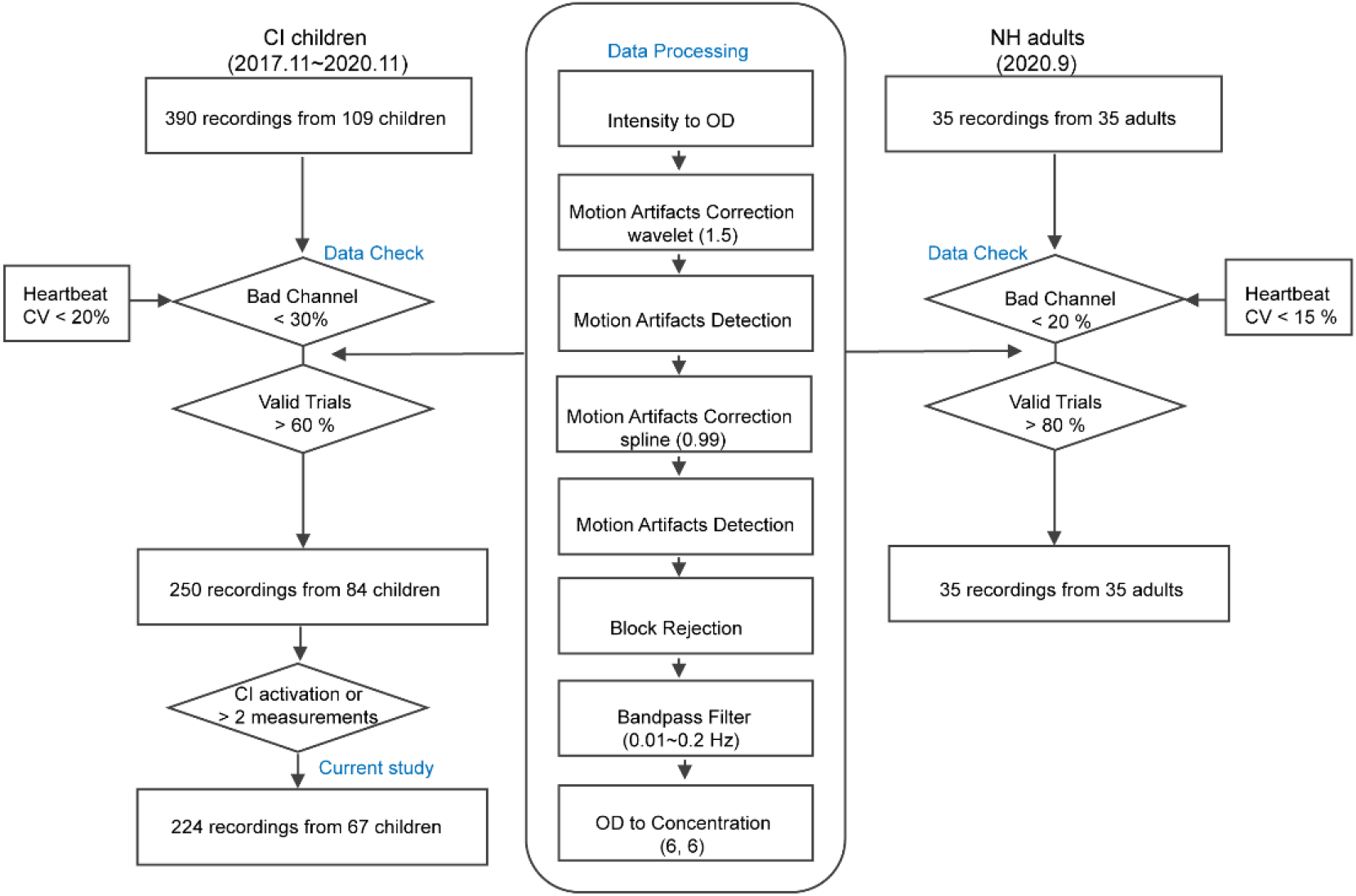
Related to Methods. Illustration of fNIRS data preprocessing and processing procedures. Abbreviations: CI, Cochlear implanted; NH, normal hearing.

## Supplemental Tables

**Table S1.**
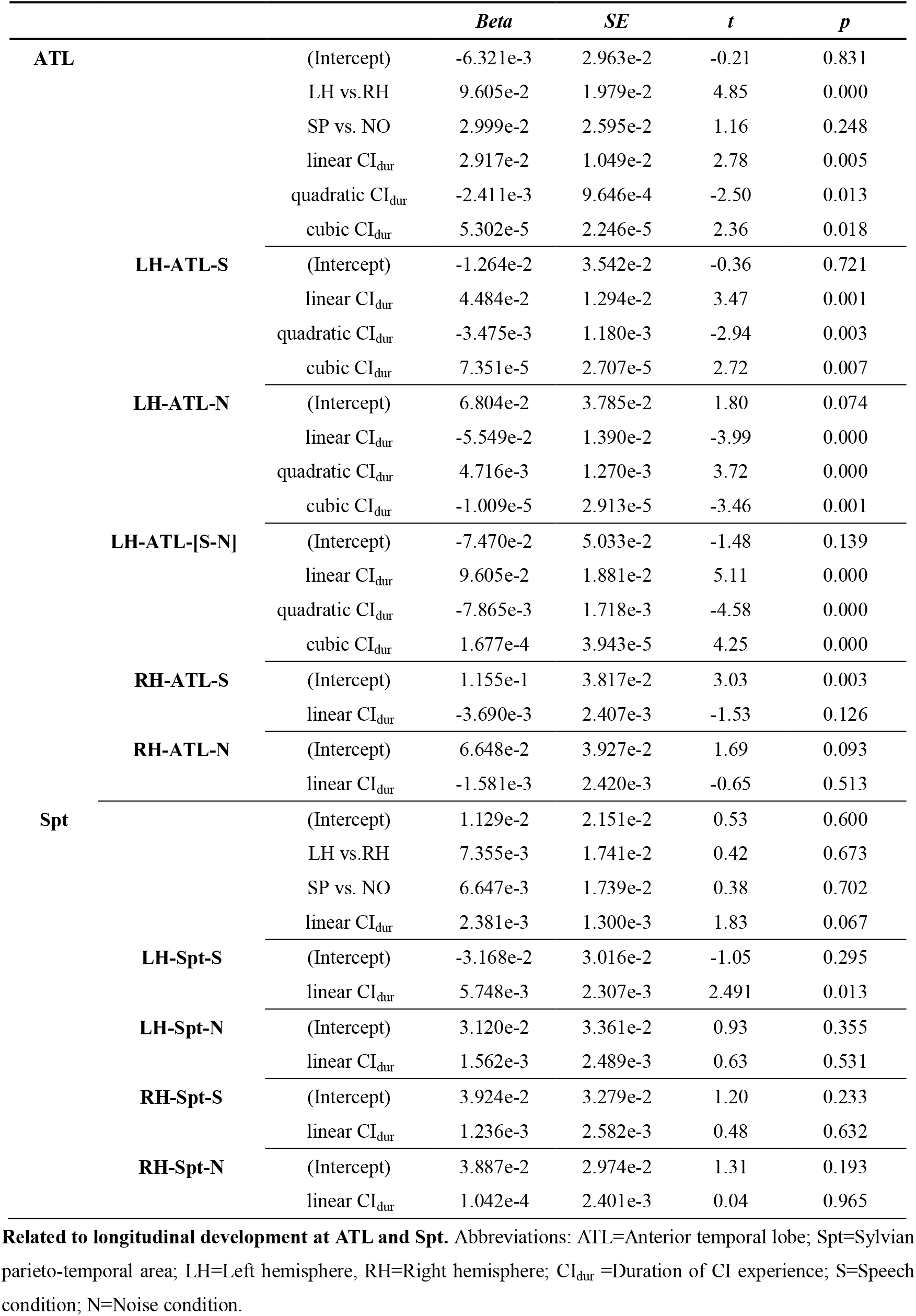
Model Estimates for ATL and Spt

**Table S2.**
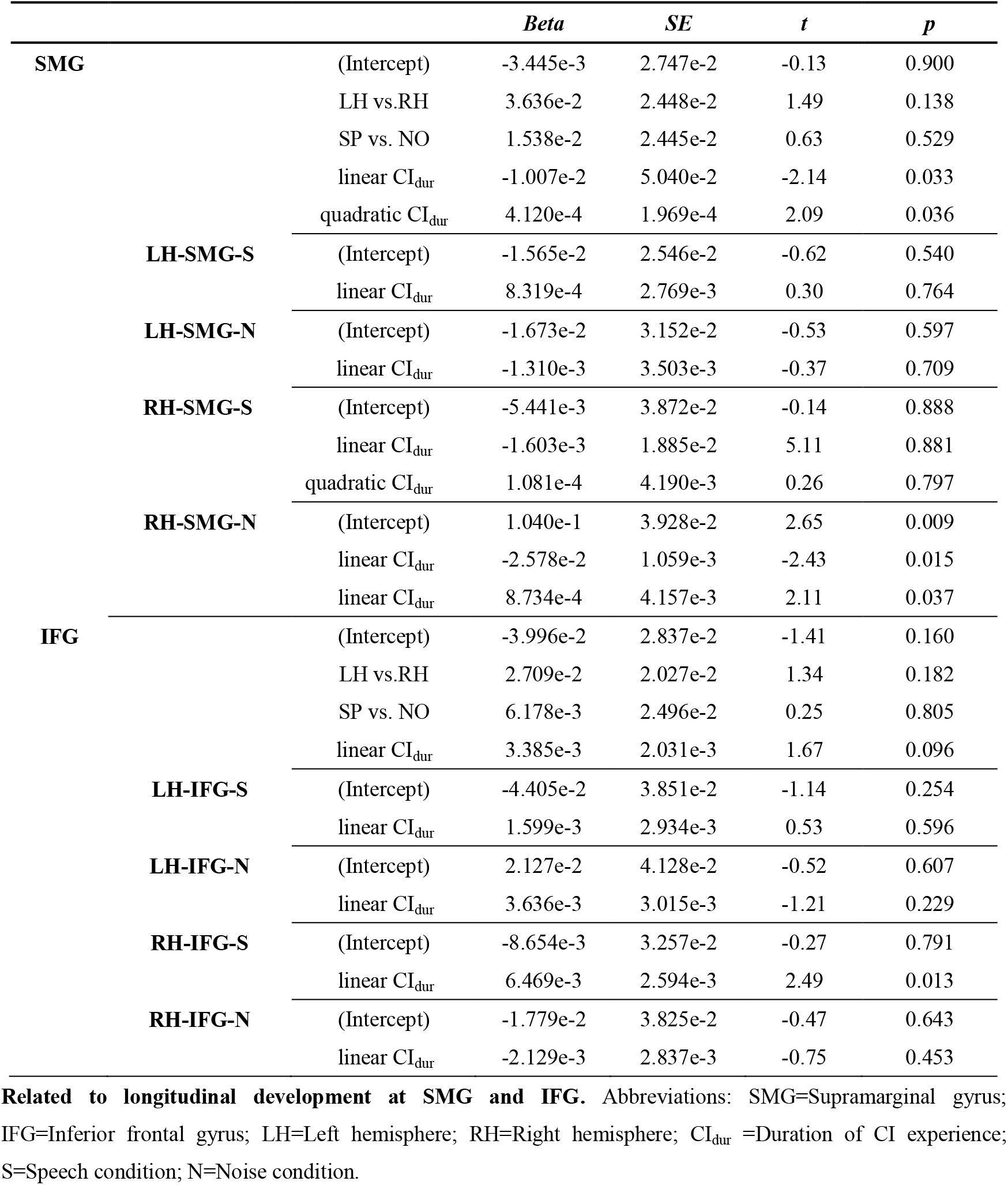
Model Estimates for SMG and IFG

**Table S3.** CI Children Demography

|  |  | VD(67) | T0(30) | DP(57) | ML(47) |
| --- | --- | --- | --- | --- | --- |
| **Gender** | female | 28 | 9 | 23 | 18 |
|  | male | 39 | 21 | 34 | 29 |
| **CI side** | Left | 24 | 10 | 20 | 18 |
|  | Right | 38 | 19 | 32 | 28 |
|  | Both | 5 | 1 | 5 | 1 |
| **CIU** | ≤90dB | 2 | 0 | 4 | 4 |
|  | 91-100dB | 23 | 10 | 20 | 18 |
|  | 101-110dB | 19 | 10 | 15 | 12 |
|  | >111dB | 21 | 10 | 16 | 13 |
| **NonCIU** | ≤90dB | 13 | 5 | 13 | 12 |
|  | 91-100dB | 17 | 8 | 13 | 11 |
|  | 101-110dB | 17 | 7 | 16 | 13 |
|  | >111dB | 18 | 10 | 13 | 11 |
| **AoI** | ≤24mo | 24 | 14 | 17 | 17 |
|  | 25-36mo | 21 | 4 | 20 | 11 |
|  | 37-48mo | 13 | 7 | 11 | 11 |
|  | >49mo | 9 | 5 | 9 | 8 |
| **HAtime** | ≤6mo | 18 | 7 | 15 | 10 |
|  | 7-12mo | 32 | 15 | 27 | 24 |
|  | 13-36mo | 12 | 5 | 11 | 10 |
|  | >37mo | 3 | 1 | 3 | 3 |

**Related to Methods.** Abbreviations: CIU = Residual hearing on CI side; NonCIU = Residual hearing on non-CI side; AoI = Age of implantation; HAtime = Duration of hearing aid; VD = Valid data; T0 = Dataset for CI activation analysis; DP = Dataset for developmental pattern analysis; ML = Dataset for machine learning analysis; mo=month.

**Table S4.**
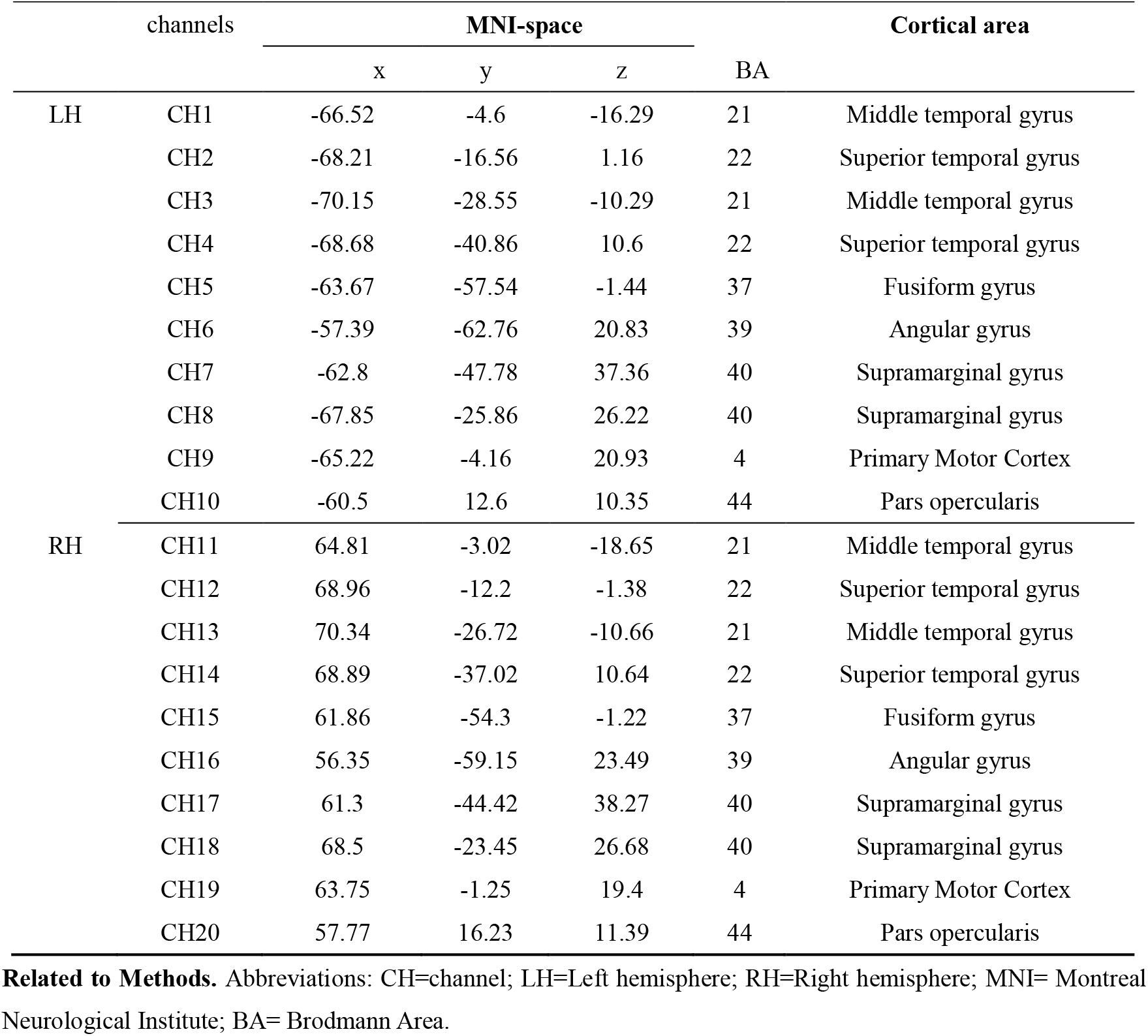
Co-registration of fNIRS channels

**Table S5.** Contrasts of LME Models for Each Region of Interest

| Areas |  | Model1 | Model2 | Model3 | Model4 | Model5 | Model6 |
| --- | --- | --- | --- | --- | --- | --- | --- |
| ATL | AIC | 2398.1 | 2398.9 | 2389.0 | 2395.4 | 2395.8 | **2385.7** |
|  | logLik | -1188.0 | -1185.5 | -1177.5 | -1184.7 | -1181.9 | **-1173.8** |
| | Contrast | A:$\chi^2$diff=5.17,p=0.159 | | B:$\chi^2$diff=15.91,p=0.001 | | C:$\chi^2$diff=21.08,p=0.002 | |
| | | D:$\chi^2$diff=5.54,p=0.136 | | E:$\chi^2$diff=16.14,p=0.001 | | F:$\chi^2$diff=21.68,p=0.001 | |
| | | G:$\chi^2$diff=6.69,p=0.035 | | H:$\chi^2$diff=7.05,p=0.029 | | I:$\chi^2$diff=7.28,p=0.026 | |
| Spt | AIC | **1230.0** | 1234.0 | 1236.5 | 1249.9 | 1229.6 | 1232.3 |
|  | logLik | **-604.99** | -604.02 | -602.25 | -612.96 | -599.81 | -598.13 |
| | Contrast | A:$\chi^2$diff=1.95,p=0.583 | | B:$\chi^2$diff=3.54,p=0.315 | | C:$\chi^2$diff=5.49,p=0.482 | |
| | | D:$\chi^2$diff=26.31,p=0.000 | | E:$\chi^2$diff=3.36,p= 0.34 | | F:$\chi^2$diff=29.66,p=0.000 | |
| | | G:$\chi^2$diff=0.00,p=1.00 | | H:$\chi^2$diff=8.42,p=0.015 | | I:$\chi^2$diff=8.24,p=0.016 | |
| SMG | AIC | 640.98 | **641.18** | 644.38 | 640.82 | 652.84 | 656.54 |
|  | logLik | -311.49 | **-308.59** | -307.19 | -309.41 | -312.42 | -311.27 |
| | Contrast | A:$\chi^2$diff=5.80,p=0.122 | | B:$\chi^2$diff=2.79,p=0.425 | | C:$\chi^2$diff=8.59,p=0.198 | |
| | | D:$\chi^2$diff=0.00,p=1.000 | | E:$\chi^2$diff=2.30,p= 0.513 | | F:$\chi^2$diff=0.00,p=1.000 | |
| | | G:$\chi^2$diff=4.16,p=0.125 | | H:$\chi^2$diff=0.00,p=1.000 | | I:$\chi^2$diff=0.00,p=1.000 | |
| IFG | AIC | **1613.0** | 1617.0 | 1620.5 | 1610.0 | 1613.6 | 1622.3 |
|  | logLik | **-796.50** | -795.53 | -794.27 | -793.02 | -791.82 | -790.13 |
| | Contrast | A:$\chi^2$diff=1.95,p=0.582 | | B:$\chi^2$diff=2.51,p=0.473 | | C:$\chi^2$diff=4.47,p=0.614 | |
| | | D:$\chi^2$diff=2.41,p=0.491 | | E:$\chi^2$diff=3.36,p=0.762 | | F:$\chi^2$diff=5.77,p=0.762 | |
| | | G:$\chi^2$diff=6.96,p=0.031 | | H:$\chi^2$diff=7.42,p=0.024 | | I:$\chi^2$diff=8.27,p=0.142 | |

**Related to Methods.** Abbreviations: ATL=Anterior temporal lobe; Spt= Sylvian parieto-temporal area; SMG=Supramarginal gyrus; IFG= Inferior frontal gyrus; LH=Left hemisphere; RH=Right hemisphere; S=Speech condition; N=Noise condition; A=Model 1 vs. Model 2; B=Model 2 vs. Model 3; C=Model 2 vs. Model3; D=Model 4 vs. Model 5; E=Model 5 vs. Model 6; F=Model 4 vs. Model 6; G=Model 1 vs. Model 4; H=Model 2 vs. Model 5; I=Model 1 vs. Model6. Optimal models were marked in bold.

**Table S6.**
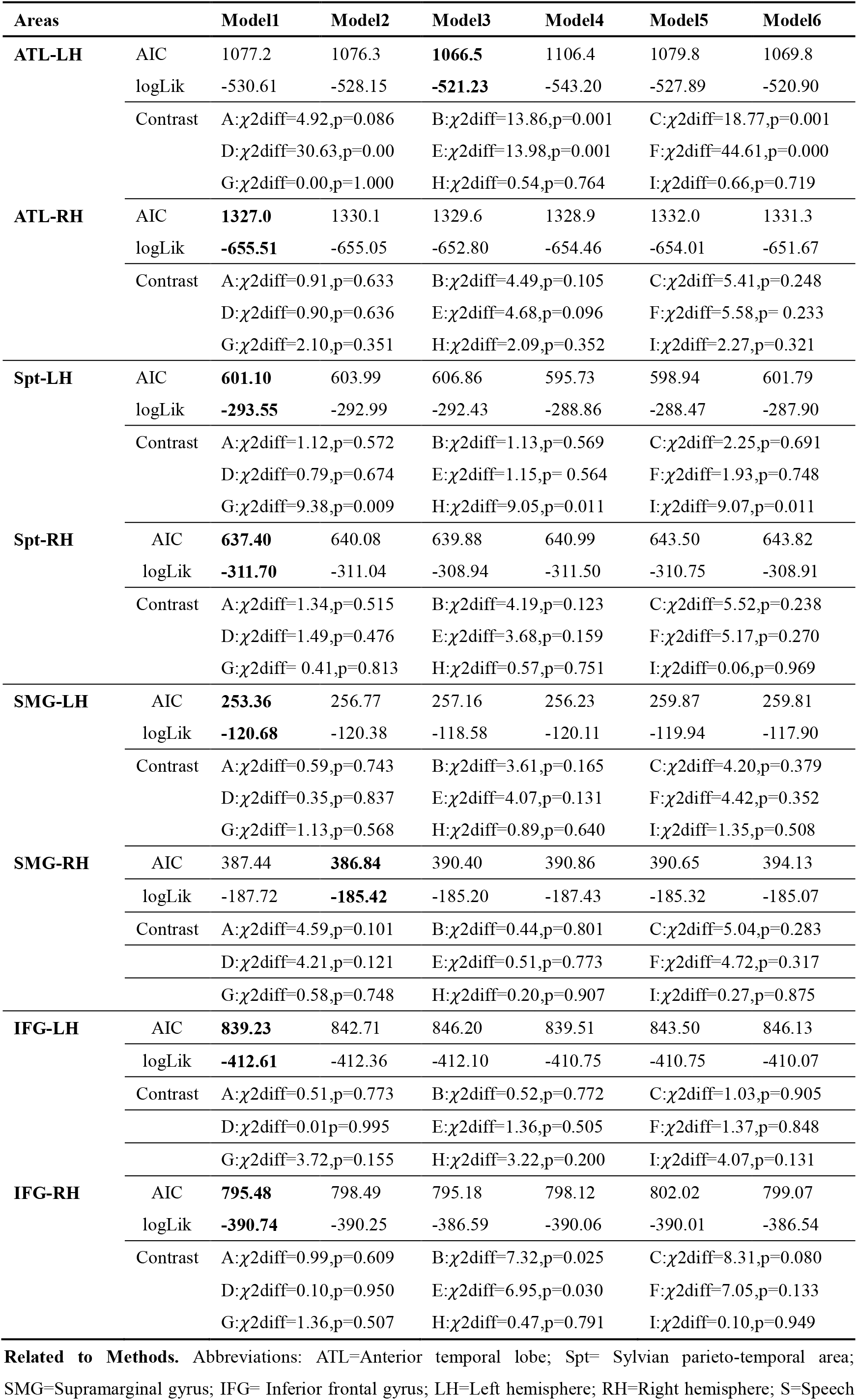
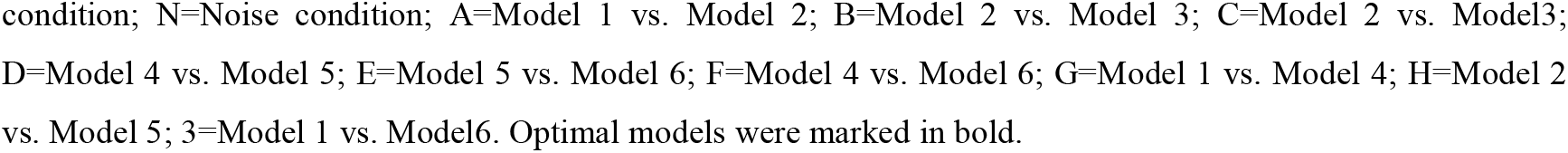
Contrasts of LME Models Separate for Each Hemisphere

| Areas |  | Model1 | Model2 | Model3 | Model4 | Model5 | Model6 |
| --- | --- | --- | --- | --- | --- | --- | --- |
| ATL-LH | AIC | 1077.2 | 1076.3 | **1066.5** | 1106.4 | 1079.8 | 1069.8 |
|  | logLik | -530.61 | -528.15 | **-521.23** | -543.20 | -527.89 | -520.90 |
| | Contrast | A:$\chi^2$diff=4.92,p=0.086 | | B:$\chi^2$diff=13.86,p=0.001 | | C:$\chi^2$diff=18.77,p=0.001 | |
| | | D:$\chi^2$diff=30.63,p=0.00 | | E:$\chi^2$diff=13.98,p=0.001 | | F:$\chi^2$diff=44.61,p=0.000 | |
| | | G:$\chi^2$diff=0.00,p=1.000 | | H:$\chi^2$diff=0.54,p=0.764 | | I:$\chi^2$diff=0.66,p=0.719 | |
| ATL-RH | AIC | **1327.0** | 1330.1 | 1329.6 | 1328.9 | 1332.0 | 1331.3 |
|  | logLik | **-655.51** | -655.05 | -652.80 | -654.46 | -654.01 | -651.67 |
| | Contrast | A:$\chi^2$diff=0.91,p=0.633 | | B:$\chi^2$diff=4.49,p=0.105 | | C:$\chi^2$diff=5.41,p=0.248 | |
| | | D:$\chi^2$diff=0.90,p=0.636 | | E:$\chi^2$diff=4.68,p=0.096 | | F:$\chi^2$diff=5.58,p= 0.233 | |
| | | G:$\chi^2$diff=2.10,p=0.351 | | H:$\chi^2$diff=2.09,p=0.352 | | I:$\chi^2$diff=2.27,p=0.321 | |
| Spt-LH | AIC | **601.10** | 603.99 | 606.86 | 595.73 | 598.94 | 601.79 |
|  | logLik | **-293.55** | -292.99 | -292.43 | -288.86 | -288.47 | -287.90 |
| | Contrast | A:$\chi^2$diff=1.12,p=0.572 | | B:$\chi^2$diff=1.13,p=0.569 | | C:$\chi^2$diff=2.25,p=0.691 | |
| | | D:$\chi^2$diff=0.79,p=0.674 | | E:$\chi^2$diff=1.15,p= 0.564 | | F:$\chi^2$diff=1.93,p=0.748 | |
| | | G:$\chi^2$diff=9.38,p=0.009 | | H:$\chi^2$diff=9.05,p=0.011 | | I:$\chi^2$diff=9.07,p=0.011 | |
| Spt-RH | AIC | **637.40** | 640.08 | 639.88 | 640.99 | 643.50 | 643.82 |
|  | logLik | **-311.70** | -311.04 | -308.94 | -311.50 | -310.75 | -308.91 |
| | Contrast | A:$\chi^2$diff=1.34,p=0.515 | | B:$\chi^2$diff=4.19,p=0.123 | | C:$\chi^2$diff=5.52,p=0.238 | |
| | | D:$\chi^2$diff=1.49,p=0.476 | | E:$\chi^2$diff=3.68,p=0.159 | | F:$\chi^2$diff=5.17,p=0.270 | |
| | | G:$\chi^2$diff= 0.41,p=0.813 | | H:$\chi^2$diff=0.57,p=0.751 | | I:$\chi^2$diff=0.06,p=0.969 | |
| SMG-LH | AIC | **253.36** | 256.77 | 257.16 | 256.23 | 259.87 | 259.81 |
|  | logLik | **-120.68** | -120.38 | -118.58 | -120.11 | -119.94 | -117.90 |
| | Contrast | A:$\chi^2$diff=0.59,p=0.743 | | B:$\chi^2$diff=3.61,p=0.165 | | C:$\chi^2$diff=4.20,p=0.379 | |
| | | D:$\chi^2$diff=0.35,p=0.837 | | E:$\chi^2$diff=4.07,p=0.131 | | F:$\chi^2$diff=4.42,p=0.352 | |
| | | G:$\chi^2$diff=1.13,p=0.568 | | H:$\chi^2$diff=0.89,p=0.640 | | I:$\chi^2$diff=1.35,p=0.508 | |
| SMG-RH | AIC | 387.44 | **386.84** | 390.40 | 390.86 | 390.65 | 394.13 |
|  | logLik | -187.72 | **-185.42** | -185.20 | -187.43 | -185.32 | -185.07 |
| | Contrast | A:$\chi^2$diff=4.59,p=0.101 | | B:$\chi^2$diff=0.44,p=0.801 | | C:$\chi^2$diff=5.04,p=0.283 | |
| | | D:$\chi^2$diff=4.21,p=0.121 | | E:$\chi^2$diff=0.51,p=0.773 | | F:$\chi^2$diff=4.72,p=0.317 | |
| | | G:$\chi^2$diff=0.58,p=0.748 | | H:$\chi^2$diff=0.20,p=0.907 | | I:$\chi^2$diff=0.27,p=0.875 | |
| IFG-LH | AIC | **839.23** | 842.71 | 846.20 | 839.51 | 843.50 | 846.13 |
|  | logLik | **-412.61** | -412.36 | -412.10 | -410.75 | -410.75 | -410.07 |
| | Contrast | A:$\chi^2$diff=0.51,p=0.773 | | B:$\chi^2$diff=0.52,p=0.772 | | C:$\chi^2$diff=1.03,p=0.905 | |
| | | D:$\chi^2$diff=0.01p=0.995 | | E:$\chi^2$diff=1.36,p=0.505 | | F:$\chi^2$diff=1.37,p=0.848 | |
| | | G:$\chi^2$diff=3.72,p=0.155 | | H:$\chi^2$diff=3.22,p=0.200 | | I:$\chi^2$diff=4.07,p=0.131 | |
| IFG-RH | AIC | **795.48** | 798.49 | 795.18 | 798.12 | 802.02 | 799.07 |
|  | logLik | **-390.74** | -390.25 | -386.59 | -390.06 | -390.01 | -386.54 |
| | Contrast | A:$\chi^2$diff=0.99,p=0.609 | | B:$\chi^2$diff=7.32,p=0.025 | | C:$\chi^2$diff=8.31,p=0.080 | |
| | | D:$\chi^2$diff=0.10,p=0.950 | | E:$\chi^2$diff=6.95,p=0.030 | | F:$\chi^2$diff=7.05,p=0.133 | |
| | | G:$\chi^2$diff=1.36,p=0.507 | | H:$\chi^2$diff=0.47,p=0.791 | | I:$\chi^2$diff=0.10,p=0.949 | |

**Related to Methods.** Abbreviations: ATL=Anterior temporal lobe; Spt= Sylvian parieto-temporal area; SMG=Supramarginal gyrus; IFG= Inferior frontal gyrus; LH=Left hemisphere; RH=Right hemisphere; S=Speech
condition; N=Noise condition; A=Model 1 vs. Model 2; B=Model 2 vs. Model 3; C=Model 2 vs. Model3; D=Model 4 vs. Model 5; E=Model 5 vs. Model 6; F=Model 4 vs. Model 6; G=Model 1 vs. Model 4; H=Model 2 vs. Model 5; 3=Model 1 vs. Model6. Optimal models were marked in bold.

**Table S7.**
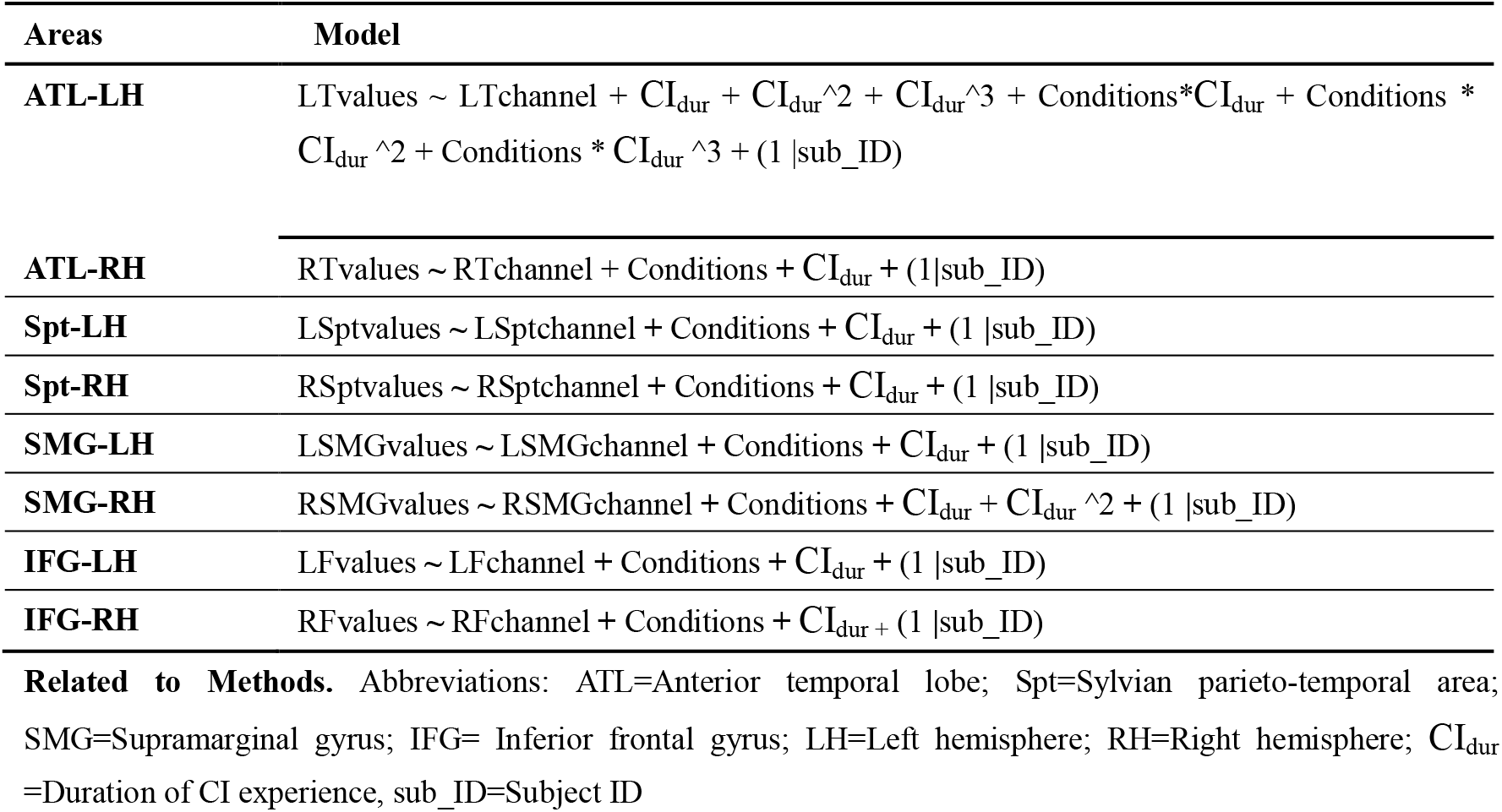
Optimal Models

## REFERENCE

Alexander-Bloch, A. F., P. T. Reiss, J. Rapoport, H. McAdams, J. N. Giedd, E. T. Bullmore and N. Gogtay (2014). “Abnormal cortical growth in schizophrenia targets normative modules of synchronized development.” Biol Psychiatry 76(6): 438–446.

Allen, M. C., T. P. Nikolopoulos and G. M. O’Donoghue (1998). “Speech Intelligibility in Children After Cochlear Implanation.” Otology & Neurotology 19(6): 742–746.

Anderson, C. A., I. M. Wiggins, P. T. Kitterick and D. Hartley (2017). “Adaptive benefit of cross-modal plasticity following cochlear implantation in deaf adults.” Proceedings of the National Academy of Sciences 114(38): 201704785.

Andics, A., J. M. Mcqueen and K. M. Petersson (2013). “Mean-based neural coding of voices.” Neuroimage 79: 351–360.

Andics, A., J. M. Mcqueen, K. M. Petersson, V. Gál and Z. Vidnyánszky (2010). “Neural mechanisms for voice recognition.” NeuroImage 52(4): 1528–1540.

Archbold, S., M. E. Lutman and D. H. Marshall (1995). “Categories of Auditory Performance.” Annals of Otology Rhinology & Laryngology Supplement 166(9): 312–314.

Arimitsu, T., M. Uchida-Ota, T. Yagihashi, S. Kojima, S. Watanabe, I. Hokuto, K. Ikeda, T. Takahashi and Y. Minagawa-Kawai (2011). “Functional Hemispheric Specialization in Processing Phonemic and Prosodic Auditory Changes in Neonates.” Frontiers in Psychology 2(202).

Baldoli, C., E. Scola, P. D. Rosa, S. Pontesilli, R. Longaretti, A. Poloniato, R. Scotti, V. Blasi, S. Cirillo and A. Iadanza (2015). “Maturation of preterm newborn brains: a fMRI–DTI study of auditory processing of linguistic stimuli and white matter development.” Brain Structure & Function 220(6): 3733.

Banich, M. T. and B. E. Depue (2015). “Recent advances in understanding neural systems that support inhibitory control.” Current Opinion in Behavioral Sciences 1: 17–22.

Belyk, Michel, Brown and Steven (2014). “Perception of affective and linguistic prosody: an ALE meta-analysis of neuroimaging studies.” Social Cognitive & Affective Neuroscience.

Bennett, I. J. and B. Rypma (2013). “Advances in functional neuroanatomy: a review of combined DTI and fMRI studies in healthy younger and older adults.” Neurosci Biobehav Rev 37(7): 1201–1210.

Bestelmeyer, P., P. Belin and M. H. Grosbras (2011). “Right temporal TMS impairs voice detection.” Current Biology 21(20): R838–R839.

Birnholz, J. C. and B. R. Benacerraf (1983). “The development of human fetal hearing.” Science 222.

Bisiacchi, P. and E. Cainelli (2021). “Structural and functional brain asymmetries in the early phases of life: a scoping review.” Brain Structure and Function(3).

Bonte, M., L. Hausfeld, W. Scharke, G. Valente and E. Formisano (2014). “Task-Dependent Decoding of Speaker and Vowel Identity from Auditory Cortical Response Patterns.” Journal of Neuroscience.

Bourke, J. D. and J. Todd (2021). “Acoustics versus linguistics? Context is Part and Parcel to lateralized processing of the parts and parcels of speech.” Laterality.

Brainard, D. H. (1997). “The Psychophysics Toolbox.” Spat Vis 10.

Brauer, J., A. Anwander, D. Perani and A. D. Friederici (2013). “Dorsal and ventral pathways in language development.” Brain & Language 127(2): 289–295.

Bulgarelli, C., C. Klerk, J. E. Richards, V. Southgate, A. Hamilton and A. Blasi (2020). “The developmental trajectory of fronto-temporoparietal connectivity as a proxy of the default mode network: a longitudinal fNIRS investigation.” Human Brain Mapping 41(10): 2717–2740.

Cabrera, L. and J. Gervain (2020). “Speech perception at birth: The brain encodes fast and slow temporal information.” Science Advances 6(30): eaba7830.

Cai, L., Q. Dong and H. Niu (2018). “The development of functional network organization in early childhood and early adolescence: A resting-state fNIRS study.” Developmental Cognitive Neuroscience 30: 223–235.

Casey, B. J., N. Tottenham, C. Liston and S. Durston (2005). “Imaging the developing brain: what have we learned about cognitive development?” Trends Cogn Sci 9(3): 104–110.

Committee, C. J. o. O. H. a. N. S. E. (2014). “Guidelines for Cochlear Implantation Work (2013).” Chinese Journal of Otorhinolaryngology Head and Neck Surgery 049(002): 89–95.

Coupe, P., G. Catheline, E. Lanuza, J. V. Manjon and I. Alzheimer’s Disease Neuroimaging (2017). “Towards a unified analysis of brain maturation and aging across the entire lifespan: A MRI analysis.” Hum Brain Mapp 38(11): 5501–5518.

Cui, X., S. Bray, D. M. Bryant, G. H. Glover and A. L. Reiss (2011). “A quantitative comparison of NIRS and fMRI across multiple cognitive tasks.” Neuroimage 54(4): 2808–2821.

David, P. B. (2008). “Praat: doing phonetics by computer (Version 5.0.45).”

Decasper, A. J. and W. P. Fifer (1980). “DeCaspar, A.J. & Fifer, W.P. Of human bonding: newborns prefer their mothers’ voice. Science 208, 1174-1176.” Science 208(4448): 1174–1176.

Dehaene-Lambertz, G., S. Dehaene and L. Hertz-Pannier (2002). “Functional neuroimaging of speech perception in infants.” Science 298(5600): 2013–2015.

Doria, V., C. F. Beckmann, T. Arichi, N. Merchant, M. Groppo, F. E. Turkheimer, S. J. Counsell, M. Murgasova, P. Aljabar, R. G. Nunes, D. J. Larkman, G. Rees and A. D. Edwards (2010). “Emergence of resting state networks in the preterm human brain.” Proceedings of the National Academy of Sciences 107(46): 20015–20020.

Eggermont, J. J. and J. K. Moore (2012). “Morphological and Functional Development of the Auditory Nervous System.” Springer Handbook of Auditory Research 42(1): 61–105.

Elia, Formisano, Federico, D. Martino, Milene, Bonte, Rainer and Goebel (2008). ““Who” is saying “what”? Brain-based decoding of human voice and speech.” Science (New York, N.Y.).

Esménio, S., J. M. Soares, P. Oliveira-Silva, Ó. F. Gonçalves, J. Decety and J. Coutinho (2019). “Brain circuits involved in understanding our own and other’s internal states in the context of romantic relationships.” Social Neuroscience.

Faust, T. E., G. Gunner and D. P. Schafer (2021). “Mechanisms governing activity-dependent synaptic pruning in the developing mammalian CNS.” Nat Rev Neurosci 22(11): 657–673.

Feng, G., E. M. Ingvalson and T. M. Griecocalub (2018). “Neural preservation underlies speech improvement from auditory deprivation in young cochlear implant recipients.” Proceedings of the National Academy of Sciences 115(5): 201717603.

Fló, A., P. Brusini, F. Macagno, M. Nespor, J. Mehler and A. L. Ferry (2019). “Newborns are sensitive to multiple cues for word segmentation in continuous speech.” Developmental Science.

Friederici, A. D. (2011). “The brain basis of language processing: from structure to function.” Physiological Reviews 91(4): 1357.

Fu, G., C. J. Mondloch, X. P. Ding, L. A. Short, L. Sun and K. Lee (2014). “The neural correlates of the face attractiveness aftereffect: A functional near-infrared spectroscopy (fNIRS) study.” Neuroimage 85: 363–371.

Gao, Wei, Alcauter, Sarael, Ramirez, Juanita, Elton, Amanda, Hernandez-Castillo and Carlos (2015). “Functional Network Development During the First Year: Relative Sequence and Socioeconomic Correlations.” Cerebral cortex 25(9): 2919–2928.

Gao, X., T. Yan, T. Huang, X. Li and Y. X. Zhang (2020). “Speech in noise perception improved by training fine auditory discrimination: far and applicable transfer of perceptual learning.” Scientific Reports 10(1).

Gavazzi, G., F. Giovannelli, T. Curro, M. Mascalchi and M. P. Viggiano (2021). “Contiguity of proactive and reactive inhibitory brain areas: a cognitive model based on ALE meta-analyses.” Brain Imaging And Behavior 15(4): 2199–2214.

Gerken, L. A. (1996). “Prosody’s role in language acquisition and adult parsing.” Journal of Psycholinguistic Research 25(2): 345–356.

Ghio, M., C. Cara and M. Tettamanti (2021). “The prenatal brain readiness for speech processing: A review on foetal development of auditory and primordial language networks.” Neuroscience And Biobehavioral Reviews 128: 709–719.

Hampshire, A., S. R. Chamberlain, M. M. Monti, J. Duncan and A. M. Owen (2010). “The role of the right inferior frontal gyrus: inhibition and attentional control.” Neuroimage 50(3): 1313–1319.

Hartwigsen, G., Y. Bengio and D. Bzdok (2021). “How does hemispheric specialization contribute to human-defining cognition?” Neuron.

Hickok, Gregory, Poeppel and David (2007). “The cortical organization of speech processing.” Nature Reviews Neuroscience.

Hickok, G. and D. Poeppel (2007). “The Cortical Organization of Speech Processing.” Nature reviews Neuroscience 8(5): 393–402.

Hoffmann, F., S. Koehne, N. Steinbeis, I. Dziobek and T. Singer (2016). “Preserved Self-other Distinction During Empathy in Autism is Linked to Network Integrity of Right Supramarginal Gyrus.” Journal of Autism and Developmental Disorders 46(2): 637–648.

Homae, F., H. Watanabe, T. Nakano and G. Taga (2007). “Prosodic processing in the developing brain.” Neuroscience Research 59(1): 29–39.

Hoshi, Y. (2007). “Functional near-infrared spectroscopy: current status and future prospects.” Journal of Biomedical Optics 12(6): 9–0.

Hua, J. Y. and S. J. Smith (2004). “Neural activity and the dynamics of central nervous system development.” Nat Neurosci 7(4): 327–332.

Huppert, T. J., S. G. Diamond, M. A. Franceschini and D. A. Boas (2009). “HomER: a review of time-series analysis methods for near-infrared spectroscopy of the brain.” Applied Optics 48(10): 0–0.

Issard, C. and J. Gervain (2018). “Variability of the hemodynamic response in infants: Influence of experimental design and stimulus complexity.” Developmental Cognitive Neuroence: 182–193.

Jöbsis, F. (1977). “Noninvasive, infrared monitoring of cerebral and myocardial oxygen sufficiency and circulatory parameters.” 198(4323): 1264.

Jusczyk, P. W., A. Cutler and N. J. Redanz (2010). “Infants’ preference for the predominant stress patterns of English words.” Child Development 64.

Klaus, B. and K. Strimmer (2015). “fdrtool: Estimation of (Local) False Discovery Rates and Higher Criticism.”

Kleiner, M. B., D. H. Brainard, D. G. Pelli, A. Ingling and C. Broussard (2007). “What’s new in Psychtoolbox-3?” Perception 36(2): 301–307.

Kral and A. (2013). “Auditory critical periods: a review from system’s perspective.” Neuroscience 247.

Kral, Hartmann, Tillein, Heid and Klinke (2002). “Hearing after congenital deafness: central auditory plasticity and sensory deprivation.” Cerebral cortex (New York, N.Y. : 1991).

Kral, A., M. F. Dorman and B. S. Wilson (2019). “Neuronal Development of Hearing and Language: Cochlear Implants and Critical Periods.” Annual Review of Neuroscience 42(1).

Kral, A., R. Hartmann, J. Tillein, S. Heid and R. Klinke (2001). “Delayed Maturation and Sensitive Periods in the Auditory Cortex.” Audiology and Neurotology 6(6): 346–362.

Kral, A., R. Hartmann, J. Tillein, S. Heid and R. Klinke (2002). “Hearing after Congenital Deafness: Central Auditory Plasticity and Sensory Deprivation.” Cerebral Cortex(8): 797.

Kral, A. and A. Sharma (2012). “Developmental neuroplasticity after cochlear implantation.” Trends in Neurosciences 35(2): 111–122.

Kral, A., J. Tillein, S. Heid, R. Klinke and R. Hartmann (2006). “Cochlear implants: cortical plasticity in congenital deprivation.” Progress in Brain Research 157: 283–313.

Kriegstein, K. V. and A. L. Giraud (2004). “Distinct functional substrates along the right superior temporal sulcus for the processing of voices.” NeuroImage 22(2): 948–955.

Kuznetsova, A., P. B. Brockhoff and R. Christensen (2016). “lmerTest: Tests in Linear Mixed Effects Models.” Journal of statistical software 2(13).

Lamm, C., J. Decety and T. Singer (2011). “Meta-analytic evidence for common and distinct neural networks associated with directly experienced pain and empathy for pain.” Neuroimage 54(3): 2492–2502.

Lana, V., T. E. Abaci, S. L. Ferradal, S. Jason, J. N. Stout, A. Banu, P. Y. Lin and G. P. Ellen (2018). “Exploring early human brain development with structural and physiological neuroimaging.” Neuroimage: S1053811918306566-.

Lebel, C. and C. Beaulieu (2011). “Longitudinal development of human brain wiring continues from childhood into adulthood.” J Neurosci 31(30): 10937–10947.

Lorenzo, R. D., L. Pirazzoli, A. Blasi, C. Bulgarelli and S. Brigadoi (2019). “Recommendations for motion correction of infant fNIRS data applicable to data sets acquired with a variety of experimental designs and acquisition systems.” NeuroImage 200.

Lu, C. M., Y. J. Zhang, B. B. Biswal, Y. F. Zang, D. L. Peng and C. Z. Zhu (2010). “Use of fNIRS to assess resting state functional connectivity.” Journal of Neuroscience Methods 186(2): 242–249.

Madhyastha, T., M. Peverill, N. Koh, C. Mccabe and K. A. Mclaughlin (2017). “Current methods and limitations for longitudinal fMRI analysis across development.” Developmental Cognitive Neuroscience 33.

Madsen, K. S., W. F. C. Baare, M. Vestergaard, A. Skimminge, L. R. Ejersbo, T. Z. Ramsoy, C. Gerlach, P. Akeson, O. B. Paulson and T. L. Jernigan (2010). “Response inhibition is associated with white matter microstructure in children.” Neuropsychologia 48(4): 854–862.

Mahmoudzadeh, M., G. Dehaene-Lambertz, M. Fournier, G. Kongolo, S. Goudjil, J. Dubois, R. Grebe and F. Wallois (2013). “Syllabic discrimination in premature human infants prior to complete formation of cortical layers.” Proceedings of the National Academy of Sciences 110(12): 4846–4851.

Malonek, D., U. Dirnagl, U. Lindauer, K. Yamada, I. Kanno and A. Grinvald (1997). “Vascular imprints of neuronal activity: Relationships between the dynamics of cortical blood flow, oxygenation, and volume changes following sensorystimulation.” Proceedings of the National Academy of Sciences of the United States of America 94(26): 14826–14831.

Martino, F. D., G. Valente, N. L. Staeren, J. Ashburner, R. Goebel and E. Formisano (2008). “Combining multivariate voxel selection and support vector machines for mapping and classification of fMRI spatial patterns.” Neuroimage 43(1): 44–58.

Meyer, M., A. Kai, A. D. Friederici, G. Lohmann and D. Cramon (2002). “FMRI reveals brain regions mediating slow prosodic modulations in spoken sentences.” Human Brain Mapping 17(2): 73–88.

Moon, M. C., Lagercrantz, H., Kuhl and K. P. (2013). “Language experienced in utero affects vowel perception after birth: A two-country study.” Acta Paediatrica International Journal of Paediatrics.

Moon, M. Christine, Fifer and P. William (2000). “Evidence of Transnatal Auditory Learning.” Journal of Perinatology.

Passingham, R. E., K. E. Stephan and R. Kotter (2002). “The anatomical basis of functional localization in the cortex.” Nat Rev Neurosci 3(8): 606–616.

Pedregosa, F., G. Varoquaux, A. Gramfort, V. Michel, B. Thirion, O. Grisel, M. Blondel, A. Müller, J. Nothman and G. Louppe (2012). “Scikit-learn: Machine Learning in Python.”

Perani, D., M. C. Saccuman, P. Scifo, D. Spada, G. Andreolli, R. Rovelli, C. Baldoli and S. Koelsch (2010). “Functional specializations for music processing in the human newborn brain.” Proceedings of the National Academy of Sciences of the United States of America 107(10).

Perani, D., M. C. Saccuman, P. Scifo, A. A. wander, D. Spada, C. Baldoli, A. Poloniato, G. Lohmann and A. D. Friederici (2011). “Neural language networks at birth.” Proceedings of the National Academy of Sciences 108(38): 16056–16061.

Piper, S. K., A. Krueger, S. P. Koch, J. Mehnert, C. Habermehl, J. Steinbrink, H. Obrig and C. H. Schmitz (2014). “A wearable multi-channel fNIRS system for brain imaging in freely moving subjects.” Neuroimage 85: 64–71.

Poeppel, D. and M. F. Assaneo (2020). “Speech rhythms and their neural foundations.” Nature reviews Neuroscience 21(Suppl. 3): 1–13.

Ralph, M., E. Jefferies, K. Patterson and T. T. Rogers (2016). “The neural and computational bases of semantic cognition.” Nature Reviews Neuroscience.

Rauschecker, J. P. and S. K. Scott (2009). “Maps and streams in the auditory cortex: nonhuman primates illuminate human speech processing.” Nature Neuroscience 12(6): 718–724.

Raznahan, A., J. P. Lerch, N. Lee, D. Greenstein, G. L. Wallace, M. Stockman, L. Clasen, P. W. Shaw and J. N. Giedd (2011). “Patterns of coordinated anatomical change in human cortical development: a longitudinal neuroimaging study of maturational coupling.” Neuron 72(5): 873–884.

Raznahan, A., P. Shaw, F. Lalonde, M. Stockman, G. L. Wallace, D. Greenstein, L. Clasen, N. Gogtay and J. N. Giedd (2011). “How does your cortex grow?” J Neurosci 31(19): 7174–7177.

Sanchez-Alonso, S. and R. N. Aslin (2020). “Predictive modeling of neurobehavioral state and trait variation across development.” Developmental Cognitive Neuroscience.

Sato, Y., Y. Sogabe and R. Mazuka (2010). “Development of Hemispheric Specialization for Lexical Pitch-Accent in Japanese Infants.” JOURNAL OF COGNITIVE NEUROSCIENCE 22(11): 2503–2513.

Schnupp, J., I. Nelken and A. King (2011). “Auditory neuroscience: Making sense of sound.”

Sevy, A., H. Bortfeld, T. J. Huppert, M. S. Beauchamp, R. E. Tonini and J. S. Oghalai (2010). “Neuroimaging with near-infrared spectroscopy demonstrates speech-evoked activity in the auditory cortex of deaf children following cochlear implantation.” Hearing Research 270(1-2): 39–47.

Shaw, P., D. Greenstein, J. Lerch, L. Clasen, R. Lenroot, N. Gogtay, A. Evans, J. Rapoport and J. Giedd (2006). “Intellectual ability and cortical development in children and adolescents.” Nature 440(7084): 676–679.

Shaw, P., N. J. Kabani, J. P. Lerch, K. Eckstrand, R. Lenroot, N. Gogtay, D. Greenstein, L. Clasen, A. Evans, J. L. Rapoport, J. N. Giedd and S. P. Wise (2008). “Neurodevelopmental trajectories of the human cerebral cortex.” J Neurosci 28(14): 3586–3594.

Shultz, S., A. Vouloumanos, R. H. Bennett and K. Pelphrey (2014). “Neural specialization for speech in the first months of life.” Developmental Science 17(5): 766–774.

Singer, T., B. Seymour, J. O’Doherty, H. Kaube, R. J. Dolan and C. D. Frith (2004). “Empathy for Pain Involves the Affective but not Sensory Components of Pain.” Science 303(5661): p.1157–1162.

Skeide, M. A. and A. D. Friederici (2016). “The ontogeny of the cortical language network.” Nature Reviews Neuroscience.

Smith, E. G., E. Condy, A. Anderson, A. Thurm and E. Redcay (2020). “Functional Near Infrared Spectroscopy in Toddlers: neural differentiation of communicative cues and relation to future language abilities.” Developmental Science.

Sowell, E. R., B. S. Peterson, P. M. Thompson, S. E. Welcome, A. L. Henkenius and A. W. Toga (2003). “Mapping cortical change across the human life span.” Nat Neurosci 6(3): 309–315.

Sowell, E. R., P. M. Thompson, C. M. Leonard, S. E. Welcome, E. Kan and A. W. Toga (2004). “Longitudinal mapping of cortical thickness and brain growth in normal children.” J Neurosci 24(38): 8223–8231.

Stiles, J. and T. L. Jernigan (2010). “The basics of brain development.” Neuropsychol Rev 20(4): 327–348.

Strangman, G., J. P. Culver, J. H. Thompson and D. A. Boas (2002). “A quantitative comparison of simultaneous BOLD fMRI and NIRS recordings during functional brain activation.” Neuroimage 17(2): 719–731.

Team, C. R. (2015). “R: A Language and Environment for Statistical Computing.” computing.

Tellis, G. and C. Tellis (2016). “Using functional near infrared spectroscopy with fluent speakers to determine haemoglobin changes in the brain during speech and non-speech tasks.” Nir News 27(3): 4.

Thatcher, R. W. (1992). “Cyclic cortical reorganization during early childhood.” Brain Cogn 20(1): 24–50.

Thatcher, R. W., D. M. North and C. J. Biver (2008). “Development of cortical connections as measured by EEG coherence and phase delays.” Hum Brain Mapp 29(12): 1400–1415.

Thatcher, R. W., R. A. Walker and S. Giudice (1987). “Human cerebral hemispheres develop at different rates and ages.” Science 236(4805): 1110–1113.

Turk, E., M. I. V. D. Heuvel, M. J. Benders, R. D. Heus and M. P. V. D. Heuvel (2019). “Functional Connectome of the Fetal Brain.” The Journal of Neuroscience 39(49): 2891–2818.

Vasung, L., E. Abaci Turk, S. L. Ferradal, J. Sutin, J. N. Stout, B. Ahtam, P. Y. Lin and P. E. Grant (2019). “Exploring early human brain development with structural and physiological neuroimaging.” Neuroimage 187: 226–254.

Velde, D., N. O. Schiller, C. C. Levelt, V. Heuven, M. Beers, J. J. Briaire and J. Frijns (2019). “Prosody perception and production by children with cochlear implants.” JOURNAL OF CHILD LANGUAGE.

Wada, S., M. Honma, Y. Masaoka, M. Yoshida and M. Izumizaki (2021). “Volume of the right supramarginal gyrus is associated with a maintenance of emotion recognition ability.” PLoS ONE 16(7): e0254623.

Walhovd, K. B., A. M. Fjell, J. Giedd, A. M. Dale and T. T. Brown (2017). “Through Thick and Thin: a Need to Reconcile Contradictory Results on Trajectories in Human Cortical Development.” Cereb Cortex 27(2): 1472–1481.

Wang, H., A. Ghaderi, X. Long, J. E. Reynolds, C. Lebel and A. B. Protzner (2021). “The longitudinal relationship between BOLD signal variability changes and white matter maturation during early childhood.” Neuroimage 242: 118448.

Wierenga, L. M., D. Van, B. Oranje, J. N. Giedd, S. Durston, J. S. Peper, T. T. Brown and E. A. Crone (2017). “A multisample study of longitudinal changes in brain network architecture in 4-13-year-old children.” Human Brain Mapping.

Xia, M., J. Wang, H. Yong and C. Peter (2013). “BrainNet Viewer: A Network Visualization Tool for Human Brain Connectomics.” Plos One 8(7): e68910.

Xu, J., X. Liu, J. Zhang, L. Zhen, X. Wang, F. Fang and H. Niu (2015). “FC-NIRS: A Functional Connectivity Analysis Tool for Near-Infrared Spectroscopy Data.” Biomed Research International 2015: 1–11.

Zhang, D., Y. Chen, X. Hou and Y. J. Wu (2019). “Near-infrared spectroscopy reveals neural perception of vocal emotions in human neonates.” Human Brain Mapping.

Zhang, D., Y. Zhou, X. Hou, Y. Cui and C. Zhou (2017). “Discrimination of emotional prosodies in human neonates: A pilot fNIRS study.” Neuroscience Letters.

Zhao, Y., X. Xiao, Y. Jiang, P. Sun, Z. Zhang, Y. Gong, Z. Li and C. Zhu (2020). “Transcranial brain atlas-based optimization for functional near-infrared spectroscopy optode arrangement: Theory, algorithm, and application.” Human Brain Mapping.

Zimmerman-Phillips, S. (1997). IT-MAIS : Infant-Toddler meaningful auditory integration scale. International Cochlear Conference.

Zong, Z. A., L. Zheng, X. F. Xiang, Z. A. Yang, D. Xnzac and E. Czad (2021). “Transcranial brain atlas for school-aged children and adolescents.” Brain Stimulation.

